# PAM18-3, a J-domain protein, maintains mitochondrial integrity and plant growth and development in *Arabidopsis thaliana*

**DOI:** 10.1101/2025.08.25.671915

**Authors:** Neha, S Souparnika, Chandan Sahi

## Abstract

Mitochondrial protein import is indispensable for organelle biogenesis and function, and is powered by the evolutionarily conserved presequence translocase-associated motor (PAM) complex. In *Arabidopsis thaliana*, three paralogs: *PAM18-1*, *PAM18-2*, and *PAM18-3*, encode J-domain proteins homologous to yeast PAM18, which stimulates the ATPase activity of mitochondrial HSP70 (mtHSP70) during protein translocation. Here, we identify *PAM18-3* as the most highly and ubiquitously expressed paralog and demonstrate its critical role in mitochondrial function and plant development. Genetic disruption of *PAM18-3* caused severe vegetative and reproductive defects, including reduced root length, smaller rosette size with fewer leaves, decreased plant height, shorter siliques, reduced seed set, and increased seed abortion. These phenotypes were fully rescued in complemented lines expressing *PAM18-3*. Ultrastructural analyses revealed profound mitochondrial abnormalities in mutants, whereas chloroplast architecture remained unaffected. Functional assays showed reduced mitochondrial membrane potential, and altered respiratory flux with a compensatory induction of the alternative oxidase (AOX) pathway. Transcript profiling revealed upregulation of AOX genes and multiple components of the mitochondrial TIM23 import apparatus and associated chaperones. Import assays demonstrated reduced mitochondrial accumulation of canonical TIM23 substrates, including IDH, ATPβ, and SHMT1, confirming a defect in matrix protein translocation. Consistently, *pam18-3* mutants accumulated elevated reactive oxygen species (ROS) and exhibited strong induction of mitochondrial dysfunction stimulon (MDS) genes, including key transcription factors mediating retrograde signaling. Together, our findings establish *PAM18-3* as a central component of the mitochondrial protein import machinery, supporting plant growth and development in *Arabidopsis thaliana*.

## Introduction

Mitochondria serve as dynamic hubs orchestrating energy production, metabolic regulation, and cellular signaling, shaping diverse facets of eukaryotic cells (Murphy, 2009; Deng et al., 2014; Faas and de Vos, 2020; Welchen et al., 2021). More than 99% of mitochondrial proteins are synthesized as precursors in the cytosol and are imported post-translationally with the assistance of cytosolic chaperones (Becker et al., 2019; Hansen and Herrmann, 2019). Additionally, evolutionarily conserved translocase complexes located in the outer and inner mitochondrial membranes, direct precursor proteins to specific mitochondrial subcompartments, including the outer and the inner membrane, intermembrane space, as well as the matrix (Endo et al., 2011; Neupert, 2015; Pfanner et al., 2019). These translocases operate within larger protein networks that integrate protein import with mitochondrial biogenesis, metabolism, and quality control (Hansen and Herrmann, 2019). Disruption of mitochondrial protein import causes disorders such as Deafness Dystonia Syndrome (DDS) and Dilated Cardiomyopathy with Ataxia in humans, and embryonic lethality, immune dysfunction, growth retardation, and sterility in plants (Davey, 2005; MacKenzie and Payne, 2007; Murcha et al., 2014; Wachoski-Dark et al., 2022).

The translocase of the outer mitochondrial membrane (TOM) complex functions as an entry gate to mitochondria, mediating the recognition and translocation of precursor proteins (Araiso and Endo, 2022; Wang et al., 2020). At the inner membrane, TIM17:23 complex forms the sole gateway for translocation of precursor proteins containing N-terminal targeting signals via the general import pathway into the matrix (Neupert and Herrmann, 2007; Chacinska et al., 2009).

The TIM17:23 complex consists of a translocation channel formed by TIM17 and TIM23 and a presequence translocase-associated motor (PAM), which drives ATP-dependent protein translocation into the matrix. The core of the PAM motor is mitochondrial HSP70 (mtHSP70), a chaperone whose activity is stimulated by J-domain proteins (JDP), PAM18 (Truscott et al., 2003; D’Silva et al., 2003; Mokranjac et al., 2003). PAM18 enhances the ATPase activity of mtHSP70 and forms a regulatory complex with PAM16, a J-like protein that modulates this stimulation, thereby fine-tuning the activity of the import motor (D’Silva et al., 2008; Frazier et al., 2004; Iosefson et al., 2007; Li et al., 2004; Pais et al., 2011).

PAM18 orthologs have been extensively studied in non-plant systems. In yeast, PAM18 is essential for protein import into the matrix and supports the assembly of respiratory complexes, particularly complex IV (D’Silva et al., 2003; Popov-Čeleketić et al., 2011; Schendzielorz et al., 2018; Truscott et al., 2003). In humans, two orthologs, *DNAJC19* and *DNAJC15*, have been identified. *DNAJC19* is essential for mitochondrial biogenesis and protein import, with mutations leading to severe disorders such as dilated cardiomyopathy with ataxia syndrome (Davey, 2005; Sinha et al., 2016). In contrast, *DNAJC15* appears to play a supportive but non-essential role in these processes (Sinha et al., 2016). However, the specific functional contributions of PAM18 paralogs to mitochondrial protein import in plants have yet to be elucidated.

In plants, the PAM components have undergone gene family expansion, resulting in functional diversification (Murcha et al., 2014). *Arabidopsis thaliana* contains two mtHSP70 paralogs (*mtHSP70-1* and *mtHSP70-2*), two PAM16 paralogs (*PAM16* and *PAM16L*), and three PAM18 paralogs (*PAM18-1*, *PAM18-2*, and *PAM18-3*) (Chen et al., 2013; Tamadaddi et al., 2021). This gene expansion has led to sub-functionalization and neo-functionalization among mitochondrial import components. For instance, knockout of *mtHSP70-1* causes severe growth defects, including reduced rosette size, shorter roots and stems, and ovule abortion (Wei et al., 2019). Additionally, *mtHSP70-1* is essential for the assembly of respiratory complex IV, maintenance of redox homeostasis, auxin-mediated embryo development, iron–sulfur (Fe–S) cluster biogenesis, and clathrin-mediated endocytosis (Li et al., 2021; Shen et al., 2025). In contrast, *mtHSP70-2* mutants exhibit no apparent developmental phenotype but play a role in Fe–S cluster assembly assembly (Zhang et al., 2025). Similarly, *PAM16* is essential for viability and negatively regulates plant immunity, while *PAM16L* contributes to early growth and stress responses (Huang et al., 2013; Tamadaddi et al., 2021). The double mutant *pam16/pam16L* is lethal, indicating overlapping but essential functions (Huang et al., 2013).

In this study, we characterized *PAM18-3*, the most abundantly expressed isoform across various tissues and developmental stages, as revealed by transcript profiling and GUS reporter assays . Analysis of *pam18-3* insertion mutant lines revealed multiple developmental abnormalities, including shortened roots, reduced rosette size, decreased leaf number, and ovule abortion. Biochemical and molecular analyses revealed elevated ROS levels, altered respiration, and de-regulated expression of genes related to mitochondrial protein import. Importantly, the efficiency of matrix-targeted precursor import via the TIM17:23 pathway was significantly reduced in the mutants. These results demonstrate that *PAM18-3* is a crucial component of the mitochondrial protein import machinery and is essential for proper mitochondrial function and normal development in *Arabidopsis thaliana*.

## Materials and methods

### Plant materials and growth conditions

*Arabidopsis thaliana* ecotype Columbia-0 (Col-0) was used as the wild-type (WT). Two independent *PAM18-3* T-DNA insertion lines, designated *pam18-3-1* (SALK_094163) and *pam18-3-2* (SALK_091892C), were obtained from the Arabidopsis Biological Resource Center (ABRC; http://abrc.osu.edu/). Seeds were surface sterilized and grown on half-strength Murashige and Skoog (½ MS) medium under controlled environmental conditions: 22 °C during the light period and 18 °C during the dark period, with a 16 h light/8 h dark photoperiod. Light intensity was maintained at 80–100 μmol m⁻² s⁻¹. For soil-grown plants, seedlings were transferred to soil after 10-14 days and maintained under the same growth conditions.

### Generation of constructs and transgenic lines

To generate the complementation construct for the *pam18-3* mutant, full-length *PAM18-3* coding sequence was amplified from *Arabidopsis thaliana* Col-0 cDNA and cloned into the pCAMBIA1300-derived vector HBP047 (Supplementary Fig. S2) under the control of the Cauliflower Mosaic Virus (CaMV) 35S promoter using Gateway cloning (Invitrogen; Catalog Nos. 12535-019 and 12535-027). For promoter analysis, approximately 2 kb of sequence upstream of the start codon of each *PAM18* gene was amplified from genomic DNA and cloned into the pGWB3 binary vector containing the *β*-glucuronidase (GUS) reporter gene. All primer sequences used for cloning are listed in Supplementary Table S1. The resulting constructs were introduced into *Agrobacterium tumefaciens* strain GV3101 and transformed into *Arabidopsis thaliana* Col-0 or *pam18-3* mutant plants using the floral dip method. Transgenic seedlings carrying the complementation construct were selected on ½ MS medium supplemented with Hygromycin B, while GUS reporter lines were selected on medium containing Kanamycin. All experiments were performed using T3 generation plants.

### RNA extraction and quantitative RT–PCR analysis

Total RNA was isolated from 10-day-old *Arabidopsis thaliana* seedlings grown on ½MS medium using TRIzol reagent (Invitrogen, USA) according to the manufacturer’s protocol. 1–2 μg of total RNA was reverse-transcribed into cDNA using the iScript™ cDNA Synthesis Kit (Bio-Rad, USA). Quantitative real-time PCR (RT-qPCR) was performed using gene-specific primers (listed in Supplementary Table S1) using the iScript SYBR Green Premix (Bio-Rad USA) on a QuantStudio™ 3 Real-Time PCR System (Applied Biosystems, Waltham, MA, USA). All reactions were run in triplicate with three independent biological replicates. Relative gene expression was normalized against two internal reference genes, and data analysis was performed using the ΔΔCt method (Livak and Schmittgen, 2001).

### GUS staining analysis

To analyse the expression patterns of PAM18 paralogs, β-glucuronidase (GUS) histochemical staining was performed using T3 generation transgenic Arabidopsis lines harboring *proPAM18-GUS* constructs. GUS staining was carried out following the protocol described by Weigel and Glazebrook (2002). After staining, the samples were mounted on a white background and imaged using an Leica DM2500 (Leica Microsystems, Wetzlar, Germany).

### Preparation of crude mitochondria from Arabidopsis seedlings

Mitochondria were isolated from *Arabidopsis thaliana* Col-0 seedlings as described by Zmudjak et al. (2017), with minor modifications. 14-day-old seedlings (200 mg), grown on MS medium, were harvested and homogenized in 2 mL of ice-cold extraction buffer containing 75 mM MOPS-KOH (pH 7.6), 0.6 M sucrose, 4 mM EDTA, 0.2% (w/v) polyvinylpyrrolidone-40, 8 mM L-cysteine, 0.2% (w/v) bovine serum albumin (BSA), and a protease inhibitor cocktail (Complete Mini, Roche Diagnostics, Germany). The homogenate was centrifuged at 1,300 × *g* for 4 min at 4 °C to remove debris. The supernatant was centrifuged at 22,000 × *g* for 50 min at 4 °C to pellet organellar membranes, including mitochondria. The pellet was washed once with 1 mL of wash buffer (37.5 mM MOPS-KOH, 0.3 M sucrose, and 2 mM EDTA, pH 7.6), followed by a second centrifugation at 22,000 × *g* for 50 min at 4 °C. The final pellet was resuspended in 200 μL of suspension buffer containing 10 mM MOPS-KOH (pH 7.2) and 0.3 M sucrose. Protein concentration was determined using the Bradford assay (Bio-Rad, USA) with BSA as a standard. For immunoblot analysis, 30 μg of mitochondrial protein was loaded per lane.

### Measurement of Oxygen uptake

Oxygen consumption was measured using a Clark-type oxygen electrode (Oxygraph, Hansatech Instruments Ltd., UK) as described by Wei et al. (2019), with minor modifications. 10-day-old *Arabidopsis thaliana* seedlings (100 mg) were pre-incubated in darkness for 30 minutes to stabilize respiration. Seedlings were then transferred into 2 mL of oxygen measurement buffer containing 300 mM mannitol, 1% (w/v) bovine serum albumin (BSA), 10 mM potassium phosphate (pH 7.2), 10 mM KCl, and 5 mM MgCl₂. Oxygen uptake was recorded at 25 °C. To distinguish between different mitochondrial respiratory pathways, 10 mM sodium azide (NaN₃) was added to inhibit cytochrome c oxidase (COX), and 10 mM salicylhydroxamic acid (SHAM) was used to inhibit alternative oxidase (AOX). Decreases in oxygen uptake following inhibitor addition were used to assess pathway-specific respiration.

### Transmission Electron Microscopy (TEM)

Leaf sections from 21-day-old *Arabidopsis thaliana* plants were fixed in 2% (v/v) glutaraldehyde and 2% (v/v) paraformaldehyde prepared in 0.1 M sodium cacodylate buffer (NaCB), pH 7.2. Samples were placed in a desiccator for 2 hours to enhance infiltration and subsequently incubated on a rotator overnight at room temperature. After fixation, samples were washed three times with 0.1 M NaCB and post-fixed with 1% (w/v) osmium tetroxide in 0.1 M NaCB for 40 minutes on ice. The tissues were then rinsed twice with NaCB and once with double-distilled water. Dehydration was carried out in a graded acetone series (40%, 60%, 80%, and 100%) on ice. Samples were then infiltrated and embedded in Agar 100 epoxy resin (Agar Scientific). Ultrathin sections (∼70 nm) were cut using an ultramicrotome and mounted on 100-mesh Cu/Pd grids (Agar Scientific). Sections were post-stained with 2% uranyl acetate followed by Reynolds’ lead citrate. Images were acquired on an FEI Morgagni 268D (FEI, Eindhoven, The Netherlands) operated at 80 keV, using a Mega View III CCD camera (Olympus-SIS).

### Mitochondrial membrane potential analysis

Protoplasts were isolated from *Arabidopsis thaliana* leaf tissue following the protocol described by Yoo et al. (2007). The protoplast suspension was stained with 5 μg/mL JC-1 dye (5,5′,6,6′-tetrachloro-1,1′,3,3′-tetraethyl-imidacarbocyanine iodide; Invitrogen, T3168) and incubated for 20 min at 25 °C in the dark. After staining, protoplasts were washed three times with W5 buffer (154 mM NaCl, 125 mM CaCl_2_, 5 mM KCl in 2 mM MES buffer, pH 5.7) to remove excess dye. Fluorescence was measured using a BD FACSCelesta Flow Cytometer. JC-1 monomers (indicative of depolarized mitochondria) were detected at 514 nm excitation and 529 nm emission, while JC-1 aggregates (indicative of polarized mitochondria) were detected at 525 nm excitation and 590 nm emission. The experiment was performed using three independent biological replicates, and each sample was analyzed in duplicate to ensure technical reproducibility.

### Detection of Reactive Oxygen Species (ROS)

#### Superoxide detection

Superoxide accumulation was assessed using nitro blue tetrazolium (NBT) staining. A 0.2% (w/v) NBT solution was prepared in a 50 mM sodium phosphate buffer (pH 7.0). 10-day-old seedlings were immersed in the staining solution and subjected to vacuum infiltration for 4 hours at room temperature. After staining, chlorophyll was removed by boiling the tissues in 90% ethanol for 10 min. The decolorized samples were stored in 70% ethanol until imaging.

### Hydrogen Peroxide detection by 3,3′-Diaminobenzidine (DAB) staining

Hydrogen peroxide accumulation was visualized using 3,3′-diaminobenzidine (DAB) staining following the method of Daudi and O’Brien (2012). A 1 mg/mL DAB solution was prepared in deionized water, and the pH was adjusted to 3.7. Ten-day-old seedlings were immersed in the DAB solution and vacuum-infiltrated for 5 min, followed by incubation at room temperature for 8 hours. Post-staining, chlorophyll was cleared by boiling the seedlings in 90% ethanol for 10 minutes. Samples were subsequently stored in a glycerol:ethanol (1:4) solution prior to imaging.

### SDS-PAGE and immunoblot analysis

Proteins were separated by SDS-PAGE and transferred onto a nitrocellulose membrane (Bio-Rad) using standard electroblotting procedures. Membranes were blocked and probed with the following primary antibodies: anti-ATP synthase subunit B (AS16 3976, Agrisera; 1:3000), anti-isocitrate dehydrogenase (IDH; AS06 203A, Agrisera; 1:2000), and anti-SHMT1 (AS23 4913, Agrisera; 1:1000). After washing, membranes were incubated with a secondary anti-rabbit antibody conjugated to Alexa Fluor™ Plus 680 (Invitrogen) at a dilution of 1:20,000. Fluorescent signal detection was performed using the Bio-Rad ChemiDoc imaging system in the 700 nm infrared fluorescence channel. All immunoblot experiments were independently repeated at least three times using biological replicates.

### Statistical analyses

All experiments were performed with at least three independent biological replicates. Statistical significance was assessed using one-way analysis of variance (ANOVA), followed by Tukey’s HSD test. Analyses were conducted using GraphPad Prism 8 software (GraphPad Software, San Diego, CA, USA). Data are presented as mean ± standard deviation (SD). Statistical significance is indicated as follows: \**p* < 0.05*, **p* < 0.01, *\*\*\*p* < 0.001, and *\*\*\*\*p* < 0.0001.

### Accession numbers

*PAM18-1* (AT2G35795), *PAM18-2* (AT3G09700), *PAM18-3* (AT5G03030), *PAM16* (AT3G59280), PAM16L (AT5G61880), *TIM17-1* (AT1G20350), *TIM17-2* (AT2G37410), *TIM17-3* (AT5G11690), *TIM23-1* (AT1G17530), *TIM23-2* (AT1G72750), *TIM23-3* (AT3G04800), *TIM44-1* (AT2G20510), *TIM44-2* (AT2G36070.1), *mtHSP70-1* (AT5G02500), *mtHSP70-2* (AT5G02490), *MGE1* (AT5G55200), *MGE2* (AT4G26780), *HSP23.5* (AT5G51440), *OM66* (AT3G50930), *ANAC013* (AT1G32870), *ANAC019* (AT1G52890), *AOX1A* (AT3G22370), *AOX1B* (AT3G22360), *AOX1C* (AT3G27620), *AOX1D* (AT1G32350), *AOX2* (AT5G64210), *UPOX1* (AT2G21640), *NBD4* (AT2G20800), *ACTIN2* (AT3G18780), *GAPDH* (AT3G26650).

## Results

### *PAM18-3* is the most abundantly expressed paralog

In *Arabidopsis thaliana*, PAM18 is encoded by three homologous genes: *PAM18-1*, *PAM18-2*, and *PAM18-3* (Tamadaddi et al., 2021; Chen et al., 2018). All three paralogs share a conserved domain architecture, comprising an N-terminal transmembrane (TM) domain, an arm region (A), and the signature J-domain at the C-terminal (Figure 1A). Structural modeling using AlphaFold confirmed that the J-domain adopts the characteristic tri-helical fold typical of JDP family proteins and contains the highly conserved HPD motif, which is essential for stimulating the ATPase activity of HSP70 chaperones (Figure 1B) (Craig et al., 2006; Verma et al., 2017). Pairwise amino acid sequence alignment showed high conservation among the three paralogs, with approximately 90% sequence similarity and 82% identity (Supplementary Figure S1), suggesting that PAM18 paralogs are structurally conserved and likely share similar molecular functions.

**Figure 1.**
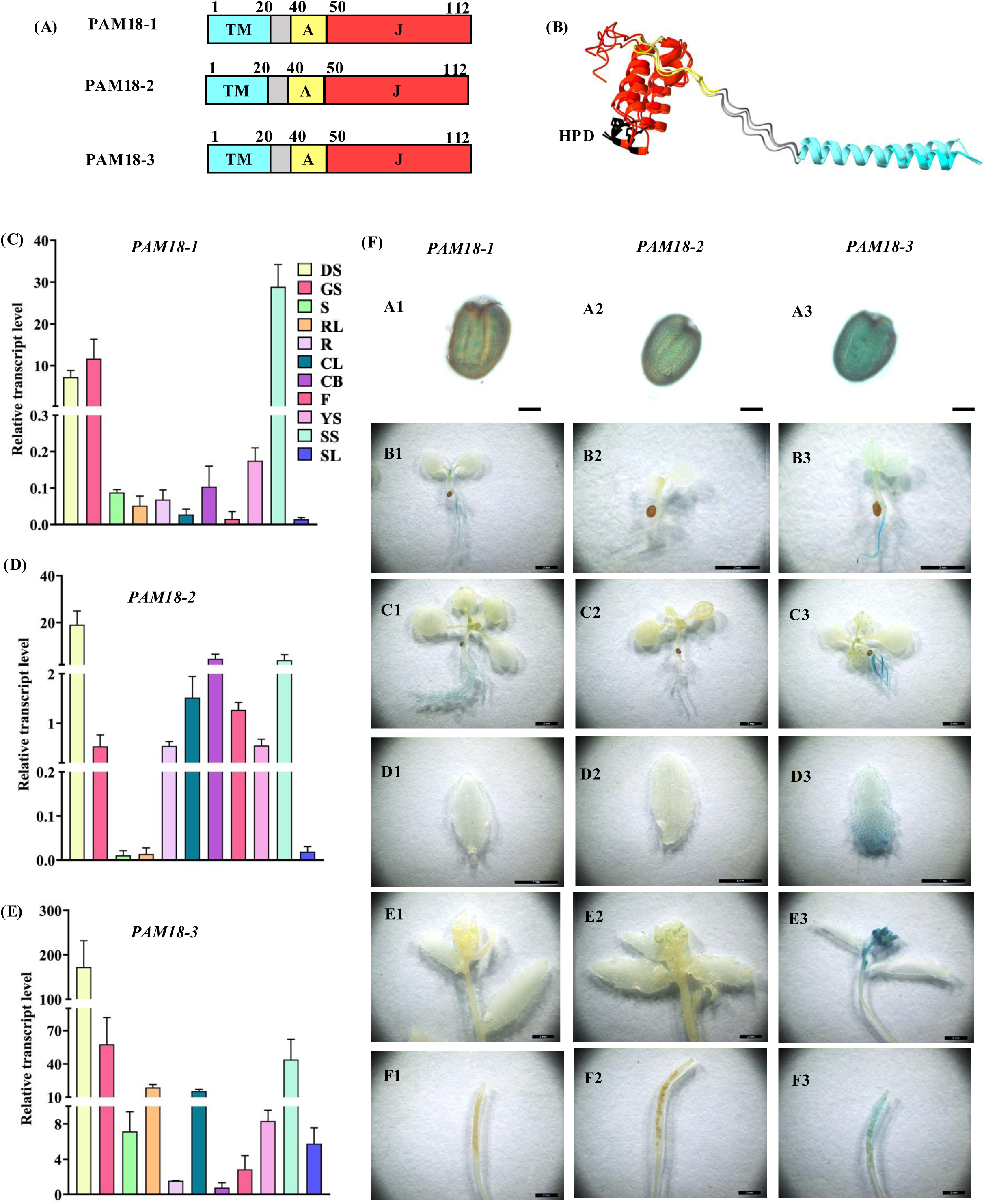
Domain architecture, structural modeling, and expression analysis of Arabidopsis PAM18 paralogs. **(A)** Schematic representation of domain organization in Arabidopsis PAM18 paralogs (*PAM18-1, PAM18-2, PAM18-3*), showing the predicted transmembrane (TM) region, arm (A), and J-domain (J). Numbers indicate amino acid positions. **(B)** Superimposition of AlphaFold-predicted structures of PAM18-1, PAM18-2, and PAM18-3, highlighting the conserved HPD motif in black. **(C–E)** Quantitative RT-PCR analysis of *PAM18-1* **(C)**, *PAM18-2* **(D)**, and *PAM18-3* **(E)** transcript levels across various Arabidopsis tissues. DS, dry seed; GS, 24 h light germinating seed; S, 7-day-old seedling; RL, rosette leaf; R, root; CL, cauline leaf; CB, closed bud; ST, stem; F, flower; YS, young silique; SS, senescent silique; SL, senescent leaf. Expression levels were normalized to *ACT2* and *GAPDH*. Data represent means ± SE of three biological replicates. **(F)** GUS reporter assay showing promoter activity of PAM18 paralogs in transgenic Arabidopsis lines: *proPAM18-1-GUS* **(A1–F1)**, *proPAM18-2-GUS* **(A2–F2)**, and *proPAM18-3-GUS* **(A3–F3)**. Staining was performed in the following tissues: A, dry seeds; B, 7-day-old seedlings; C, 14-day-old seedlings; D, rosette leaf; E, flower; F, silique. Scale bars A= 100 µm, B, C, E, F=2mm, and D=5mm.

To investigate the potential functional divergence among the Arabidopsis PAM18 paralogs, we first analyzed their expression profiles using qRT-PCR across various developmental stages and tissues. The three genes exhibited distinct expression patterns. *PAM18-1* and *PAM18-2* showed relatively low transcript levels across most developmental stages and tissues, whereas *PAM18-3* exhibited the highest expression and was expressed in nearly all tissues examined (Figure 1C-E). To further study the spatial expression patterns, stable transgenic Arabidopsis lines expressing the β-glucuronidase (GUS) reporter gene under the control of a 2 kb promoter region upstream of each paralog’s start codon were generated. Histochemical GUS staining supported our qRT-PCR findings, with *PAM18-3* showing strong and widespread GUS activity throughout the plant (Figure 1F). While *PAM18-1* was predominantly expressed in seeds and seedlings, *PAM18-2* was mainly active in seeds and floral tissues (Figure 1F). Three independent transgenic lines were analyzed for each construct, all of which displayed consistent and reproducible expression profiles. Together, these findings indicate that while the Arabidopsis PAM18 paralogs are structurally conserved, they display distinct and non-overlapping expression profiles. Among them, *PAM18-3* is the most highly and ubiquitously expressed paralog, implying a predominant role in plant development and mitochondrial protein homeostasis.

### *PAM18-3* is essential for vegetative and reproductive growth

To investigate the functional role of *PAM18-3*, two independent T-DNA insertion alleles, *pam18-3-1* (SALK_094163) and *pam18-3-2* (SALK_091892C), carrying insertions in the promoter region and third intron, respectively (Figure 2A), were identified and characterized. In parallel, two independent complemented lines (Comp 1 and Comp 2) were generated by expressing the *PAM18-3* coding sequence under the CaMV 35S promoter in the *pam18-3-1* background (Supplementary Figure S2A, B). qRT-PCR analysis confirmed that both mutant lines exhibited a strong reduction in *PAM18-3* transcript levels, with expression reduced to <10% of WT levels, while expression was restored in the complemented lines (Supplementary Figure S2B).

**Figure 2.**
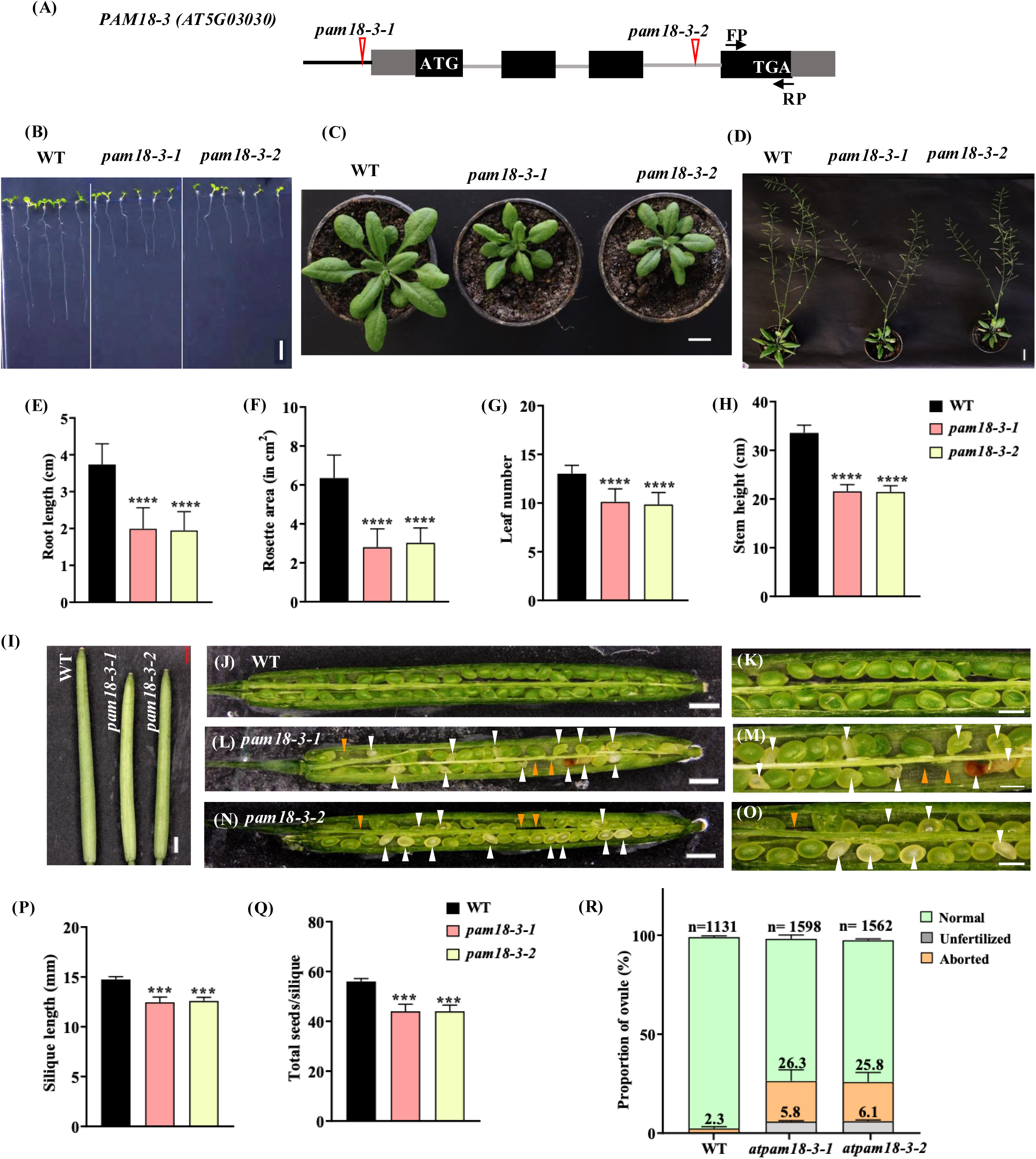
Phenotypic characterization of *pam18-3* mutants. **(A)** Schematic representation of the *PAM18-3* gene *(AT5G03030)* structure showing T-DNA insertion sites in *pam18-3-1* and *pam18-3-2* mutants (red arrowheads). Exons (black boxes), introns (gray lines), UTRs (dark gray boxes), and promoter region (black line) are indicated. Positions of primers used for RT-qPCR (FP and RP) are shown as arrows. **(B–H)** Growth phenotypes of *pam18-3* mutants. Representative images of 10-day-old seedlings showing root growth **(B)**, 21-day-old rosettes **(C)**, and 45-day-old flowering plants **(D)** in WT, *pam18-3-1*, and *pam18-3-2* lines. Quantification of root length **(E)**, rosette area **(F)**, total leaf number **(G)**, and stem height **(H)**. Data represent mean ± SD from three independent experiments (*n* = 30 plants per genotype). Statistical significance was assessed using one-way ANOVA with Tukey’s HSD test (\*\*\*\**p* < 0.0001). Scale bars: B, C = 1 cm, and D = 2 cm. **(I–R)** Reproductive phenotypes of *atpam18-3* mutants. Representative images of mature siliques at 8 days post anthesis (DPA) **(I)**. Seed development in 8-DPA siliques of WT **(J, K)**, *pam18-3-1* **(L, M)**, and *pam18-3-2* **(N, O)**. Aborted ovules are marked with white arrowheads; unfertilized ovules with orange arrowheads. Quantification of silique length **(P)**, total seeds per silique **(Q)**, and proportion of normal, unfertilized, and aborted ovules per silique **(R)**. Data represent mean ± SD from three biological replicates (*n* = 10 siliques from 10 individual plants per replicate). Statistical analysis was performed using one-way ANOVA with Tukey’s HSD test (*\*\*\*p < 0.001*). Scale bars: I, J, L, N = 1 mm, and K, M, O = 0.5 mm.

Phenotypic characterization revealed pronounced defects in both vegetative and reproductive development of *pam18-3* mutants. Rosette area, primary root length, and plant height were all significantly reduced compared to WT (Figures 2B–D). Under standard growth conditions, primary roots of 10-day-old *pam18-3-1* and *pam18-3-2* seedlings were significantly shorter, averaging ∼2.0 cm compared with ∼3.8 cm in WT (Figure 2E). At 21 days, mutants developed smaller rosettes with average areas of ∼2.4 cm² compared with ∼6.0 cm² in WT (Figure 2F), and produced fewer leaves, averaging ∼9 in mutants compared with ∼13 in WT (Figure 2G). By 45 days, adult *pam18-3* plants exhibited reduced height as well, averaging ∼21 cm compared to ∼33 cm in WT (Figure 2H).

Reproductive development was also strongly impaired, with *pam18-3* siliques averaging ∼13 mm in length compared to ∼15 mm in WT (Figure 2I, P). In addition, seed development was significantly compromised, as mutant siliques contained unfertilized ovules and wrinkled white or yellow aborted seeds (Figure 2L-O), in contrast to the uniform seed set observed in WT (Figure 2J, K). On an average, WT siliques produced ∼55 seeds, whereas *pam18-3-1* and *pam18-3-2* siliques contained ∼40 and 42 seeds, respectively (Figure 2Q). Among these, ∼26% of seeds in *pam18-3-1* and ∼28.7% in *pam18-3-2* were aborted (Figure 2R). Consistently, siliques from *pam18-3* mutants displayed elevated rates of unfertilized ovules (5–6%) relative to <1% in WT (Figure 2R). Collectively, these analyses revealed an overall ∼15% reduction in total seed set and a ∼25% increase in seed abortion in *pam18-3* compared to WT. All mutant phenotypes, including reduced rosette growth, shorter roots and stems, shorter siliques, and elevated seed abortion, were fully rescued in both the complemented lines (Supplementary Figure S2), confirming that the developmental defects were specifically attributable to disruption of *PAM18-3*. Together, these results demonstrate that *PAM18-3* is critical for vegetative growth and seed development in Arabidopsis.

### Mitochondrial ultrastructure is severely affected in *pam18-3-1* mutant plants

A previous study showed that PAM18-3 is localized to both mitochondria and chloroplasts (Gill-Hille et al., 2022; Tamadaddi et al., 2021). Given the phenotypic abnormalities observed in *pam18-3-1* mutants, we asked if loss of *PAM18-3* affected ultrastructure of mitochondria or chloroplasts. For this a comparative ultrastructural analysis of these organelles was performed using transmission electron microscopy (TEM). In wild-type (WT) leaf cells, mitochondria predominantly exhibited elongated, tubular morphologies (Figure 3A, C), consistent with healthy and metabolically active organelles (Galloway and Yoon, 2013; Karbowski and Youle, 2003; Rambold et al., 2011). In contrast, mitochondria in *pam18-3-1* mutants showed pronounced morphological abnormalities. These included rounded, swollen, and irregularly shaped organelles (Figure 3B, D), features commonly associated with mitochondrial stress and dysfunction (Galloway and Yoon, 2013; Javadov et al., 2018). Parallel TEM analysis of chloroplasts revealed no observable structural differences between WT and *pam18-3-1* mutant cells. The thylakoid membranes, grana stacking, and overall chloroplast architecture remained intact and comparable across genotypes (Figure 3E, F). These findings indicate that the primary subcellular defect in *pam18-3* mutants is mitochondrial, underscoring the crucial role of *PAM18-3* in maintaining mitochondrial integrity.

**Figure 3.**
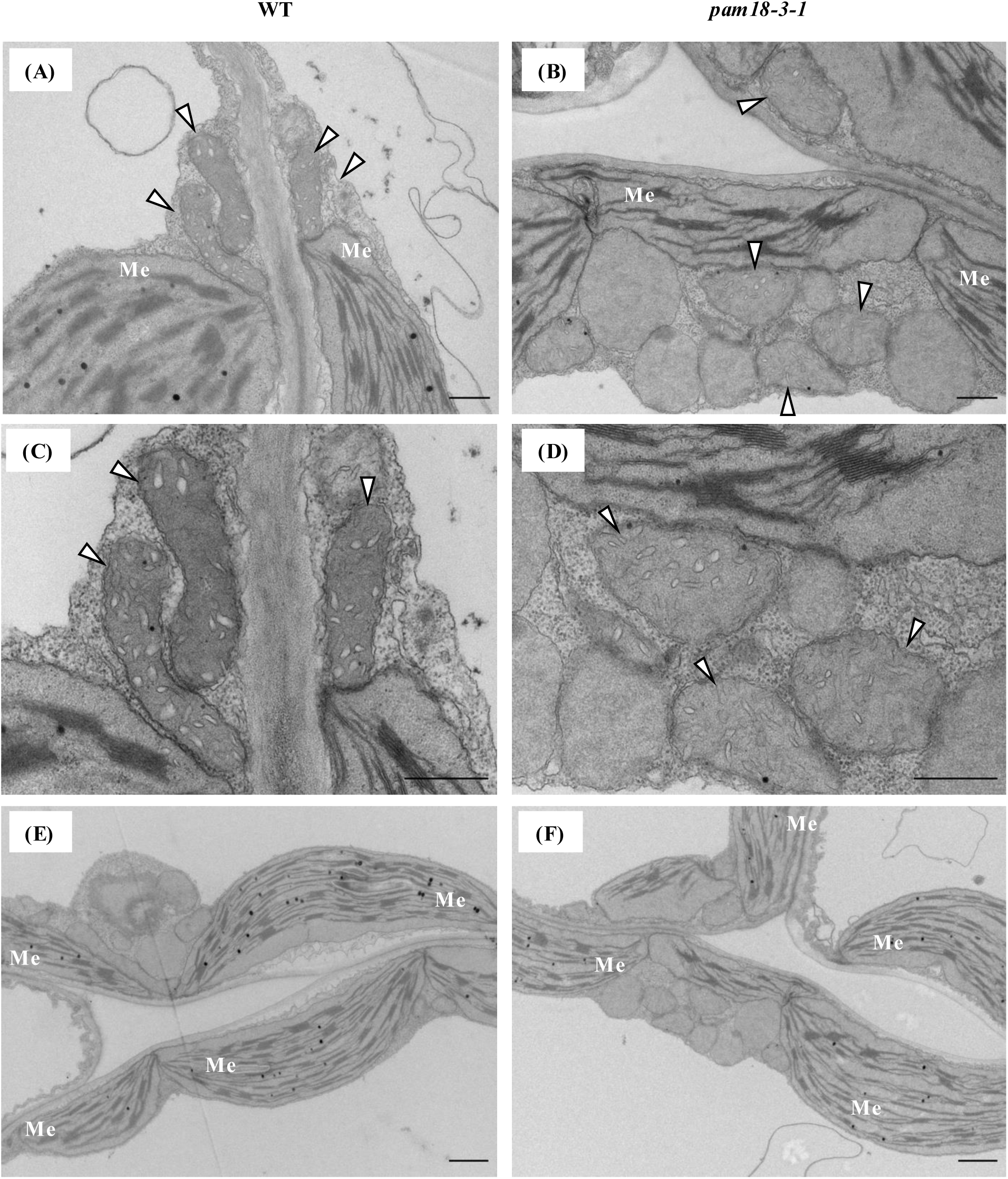
Ultrastructural analysis of mitochondria and chloroplasts in WT and *pam18-3-1* leaves. Transmission electron micrographs of leaf mesophyll cells from 21-day-old WT **(A)**, **(C)**, **(E)** and *pam18-3-1* plants **(B)**, **(D), (F)**. **(A)** Mitochondria with normal ultrastructure (indicated by arrows) adjacent to mesophyll cells (Me) in the wild type. **(B)** Mitochondria with altered morphology (arrows) in *pam18-3-1* mesophyll cells. High-magnification image showing elongated and tubular mitochondria in the WT **(C),** and irregularly shaped, swollen mitochondria in *pam18-3-1* **(D)**. **(E)** Chloroplasts in mesophyll cells of the WT. **(F)** Chloroplasts in mesophyll cells of *pam18-3-1*. Scale bars: A–D = 500 nm; E, F = 1000 nm. Images are representative of three biological replicates; 25 mitochondria were analyzed from WT and 22 from *pam18-3-1*.

### *pam18-3* mutant exhibits reduced mitochondrial membrane potential and altered respiratory activity

Mitochondrial ultrastructural abnormalities observed in *pam18-3* mutants, suggested potential impairments in mitochondrial function. To explore its physiological consequences, mitochondrial membrane potential was evaluated using the cationic dye JC-1, which fluoresces differently based on membrane polarization (Salvioli et al., 1997). In functional mitochondria, JC-1 accumulates and forms red-fluorescent aggregates, whereas depolarized mitochondria retain green-fluorescent monomers. Protoplasts isolated from leaves of wild-type (WT), *pam18-3-1*, and the complemented line (Comp 1) were stained with JC-1 and analyzed by flow cytometry. Compared to the WT, a higher proportion of green fluorescence in *pam18-3-1* protoplasts indicated reduced membrane potential and compromised mitochondrial functionality (Figure 4 A, B). This defect was rescued in Comp 1, supporting the specific role of *PAM18-3* in maintaining mitochondrial membrane potential (Figure 4C). Quantitative analysis of the red-to-green fluorescence ratios further substantiated these observations, with *pam18-3-1* showing a significantly lower ratio compared to WT and Comp 1 (Figure 4D). In addition, a fraction of cells displayed dual red and green fluorescence, representing heterogeneous mitochondrial polarization states within individual protoplasts (Supplementary Figure S4).

**Figure 4.**
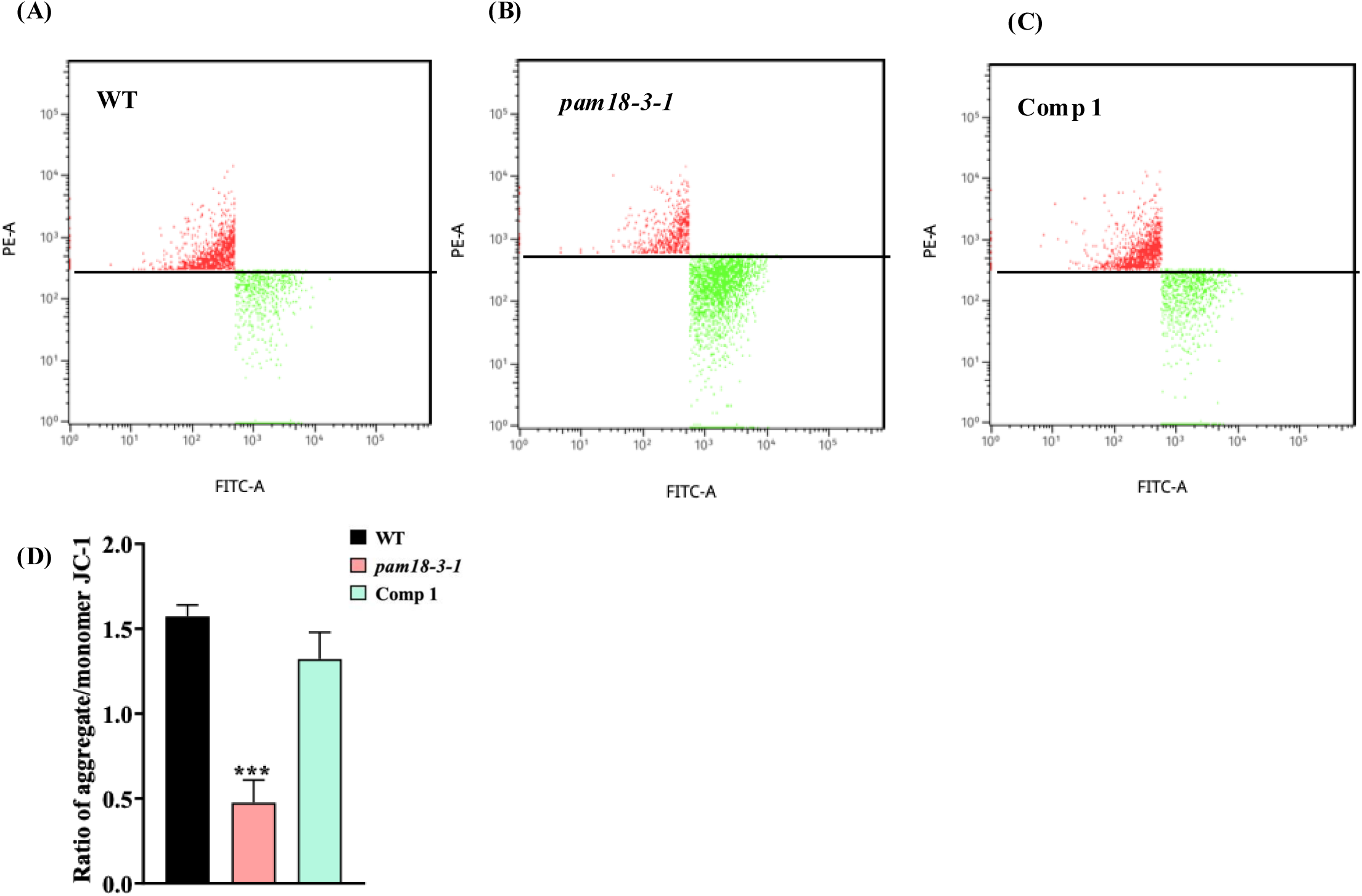
Flow cytometry analysis of mitochondrial membrane potential in Arabidopsis protoplasts. **(A–C)** Representative dot plots of JC-1-stained protoplasts isolated from wild type **(A)**, *pam18-3-1* mutant **(B)**, and complemented line Comp 1 **(C)**. The Y-axis (PE-A) represents JC-1 aggregates (red fluorescence), reflecting polarized mitochondria with intact membrane potential, while the X-axis (FITC) represents JC-1 monomers (green fluorescence), indicative of depolarized mitochondria. **(D)** Quantification of the JC-1 aggregate-to-monomer fluorescence ratio in WT, *pam18-3-1*, and Comp 1. Data represent mean ± SE from three independent experiments. Asterisks denote statistically significant differences compared to WT (one-way ANOVA with Tukey’s HSD test; \*\*\**p* < 0.001).

To further assess mitochondrial function, respiratory activity was measured in 10-day-old seedlings of WT, *pam18-3-1*, and the Comp 1, using a Clark-type oxygen electrode. Notably, under normal conditions, *pam18-3-1* seedlings exhibited a significantly elevated rate of oxygen consumption relative to WT, indicating altered respiration (Figure 5A). To investigate the basis of this elevated respiration, we analyzed the contributions of the two major mitochondrial respiratory pathways: cytochrome oxidase (COX) and alternative oxidase (AOX). Oxygen uptake was measured in the presence of specific inhibitors to distinguish the individual contributions of each of these pathways. Treatment with 10 mM sodium azide (NaN₃), a COX inhibitor, reduced oxygen consumption by approximately 63% in WT but only by 30% in *pam18-3-1* (Figure 5A). Conversely, treatment with 10 mM salicylhydroxamic acid (SHAM), an AOX inhibitor, led to a 61% reduction in oxygen uptake in *pam18-3-1* compared to a 29% reduction in WT (Figure 5A), indicating a compensatory upregulation of AOX activity in the mutant background. As expected, when both inhibitors were applied simultaneously, oxygen consumption dropped to minimal basal levels in all the genotypes (Figure 5A).

**Figure 5.**
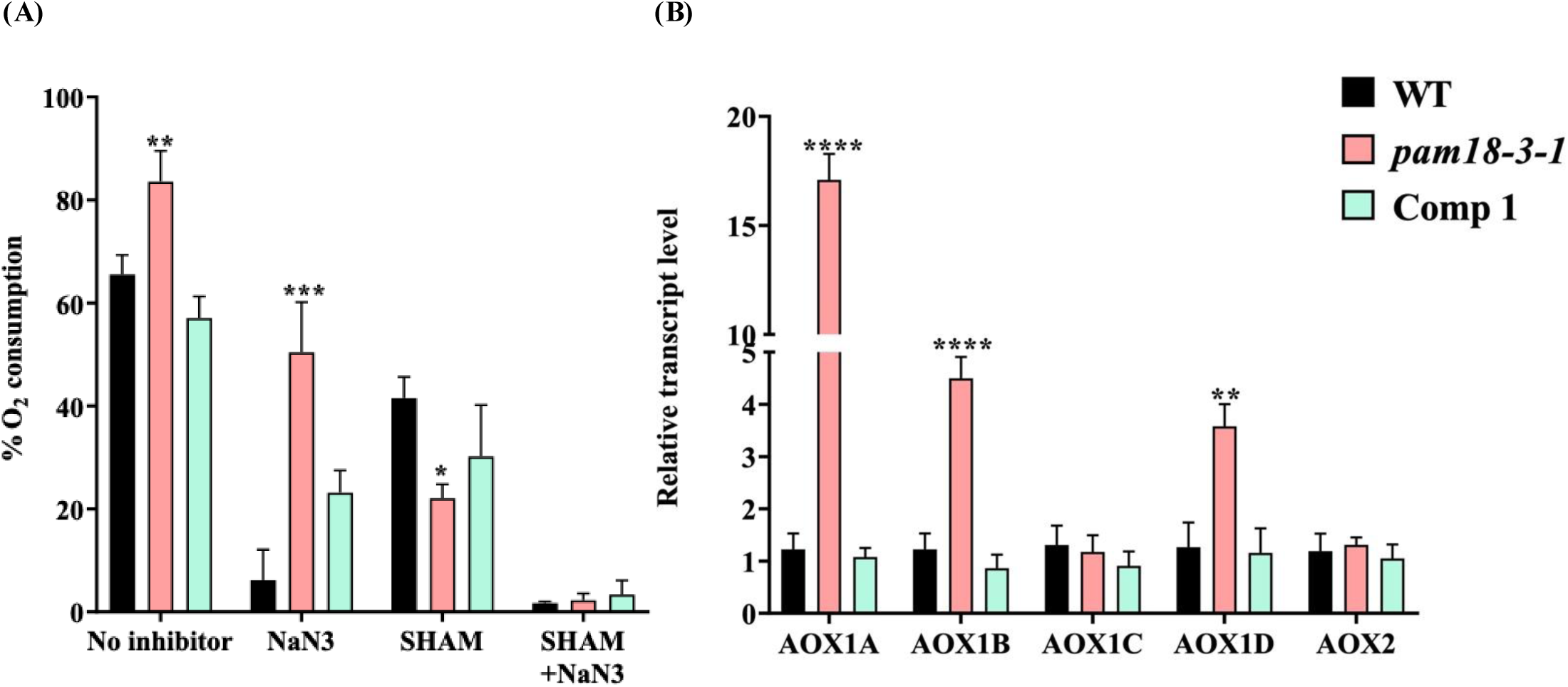
Analysis of oxygen consumption and AOX gene expression in WT, *pam18-3-1*, and complemented (Comp 1) seedlings. **(A)** Oxygen consumption in 10-day-old seedlings of wild type (WT), *pam18-3-1*, and Comp 1 under control (no inhibitor) conditions or in the presence of 10 mM sodium azide (NaN₃), 10 mM salicylhydroxamic acid (SHAM), or both. Data are shown as percentage of total O₂ consumption and represent mean ± SD of three biological replicates. Asterisks indicate significant differences from WT under the same condition (one way ANOVA with Tukey’s HSD test; \**p* < 0.05, \*\**p* < 0.01, \*\*\**p* < 0.001). **(B)** Transcript abundance of AOX genes (*AOX1A, AOX1B, AOX1C, AOX1D,* and *AOX2*) in WT, *pam18-3-1*, and Comp 1 seedlings determined by RT-qPCR. Expression was normalized to *ACT2* and *GAPDH*, with WT set to 1. Data represent mean ± SD of three biological replicates. Asterisks indicate significant differences compared to WT (one-way ANOVA with Tukey’s HSD test; \*\**p* < 0.01, \*\*\*\**p* < 0.0001).

Since *pam18-3-1* mutants displayed stronger SHAM sensitivity than WT, we hypothesized that the AOX pathway is transcriptionally upregulated in the mutants. To test this, transcript levels of five AOX genes (*AOX1A*, *AOX1B*, *AOX1C*, *AOX1D*, and *AOX2*) were quantified by qRT-PCR. Expression of *AOX1A*, *AOX1B*, and *AOX1D* was significantly upregulated in *pam18-3-1* compared to WT, whereas *AOX1C* and *AOX2* expression remained unchanged (Figure 5B). In the complemented line Comp1, both respiratory activity and AOX gene expression were restored to WT levels (Figure 5A, B), confirming that the respiratory alterations observed in *pam18-3-1* are specifically due to the loss of *PAM18-3* function. Taken together, these findings demonstrate that the absence of *PAM18-3* leads to reduced mitochondrial membrane potential, and activation of a compensatory AOX-dependent electron transport pathway to maintain mitochondrial function.

### Expression of genes encoding components of the mitochondrial TIM23 import complex is altered in the *pam18-3* mutant plants

PAM18 is a conserved co-chaperone of the TIM23-associated import motor across eukaryotes, where it plays a critical role in driving the translocation of presequence-containing proteins into the mitochondrial matrix (Mokranjac et al., 2006; Truscott et al., 2003). In both yeast and mammals, disruption of PAM18 function leads to compromised mitochondrial protein import and subsequent mitochondrial dysfunction (Sinha et al., 2016; Truscott et al., 2003). We therefore hypothesized that disruption of *PAM18-3* in *Arabidopsis thaliana* could profoundly impact the expression and coordination of the TIM23 import machinery. To test this, we examined the expression profiles of key TIM23 pathway genes in 10-day-old seedlings of WT, *pam18-3-1* mutants, and complemented (Comp 1) plants using qRT-PCR. Significant alterations in the transcript levels of multiple TIM23 pathway-related genes were observed in the *pam18-3-1* mutant background. Genes encoding the core components of the inner membrane translocon, including *TIM17-3, TIM23-1, TIM23-2,* and *TIM23-3*, were markedly upregulated in the *pam18-3-1* mutant relative to WT (Figure 6C-F). Components of the presequence translocase-associated motor (PAM) complex also showed altered expression. Notably, transcript levels of *mtHSP70-2*, encoding the mitochondrial HSP70 chaperone that facilitates ATP-dependent preprotein translocation, were significantly elevated (Figure 6N). Similarly, expression of its nucleotide exchange factors *MGE1* and *MGE2* was also increased (Figure 5O, P). In contrast, the expression of *PAM16*, which forms a regulatory heterodimer with PAM18 to modulate mtHSP70’s ATPase activity, was significantly downregulated in the mutant (Figure 6K, L). Collectively, these results suggest that the loss of *PAM18-3* perturbs mitochondrial protein import and triggers a transcriptional response possibly aimed at restoring import efficiency through upregulation of TIM23 complex components and associated chaperones. These transcriptional alterations in *pam18-3-1* were largely restored in the Comp 1, and were comparable to WT (Figure 6), confirming that the defects stem from the loss of *PAM18-3* function.

**Figure 6.**
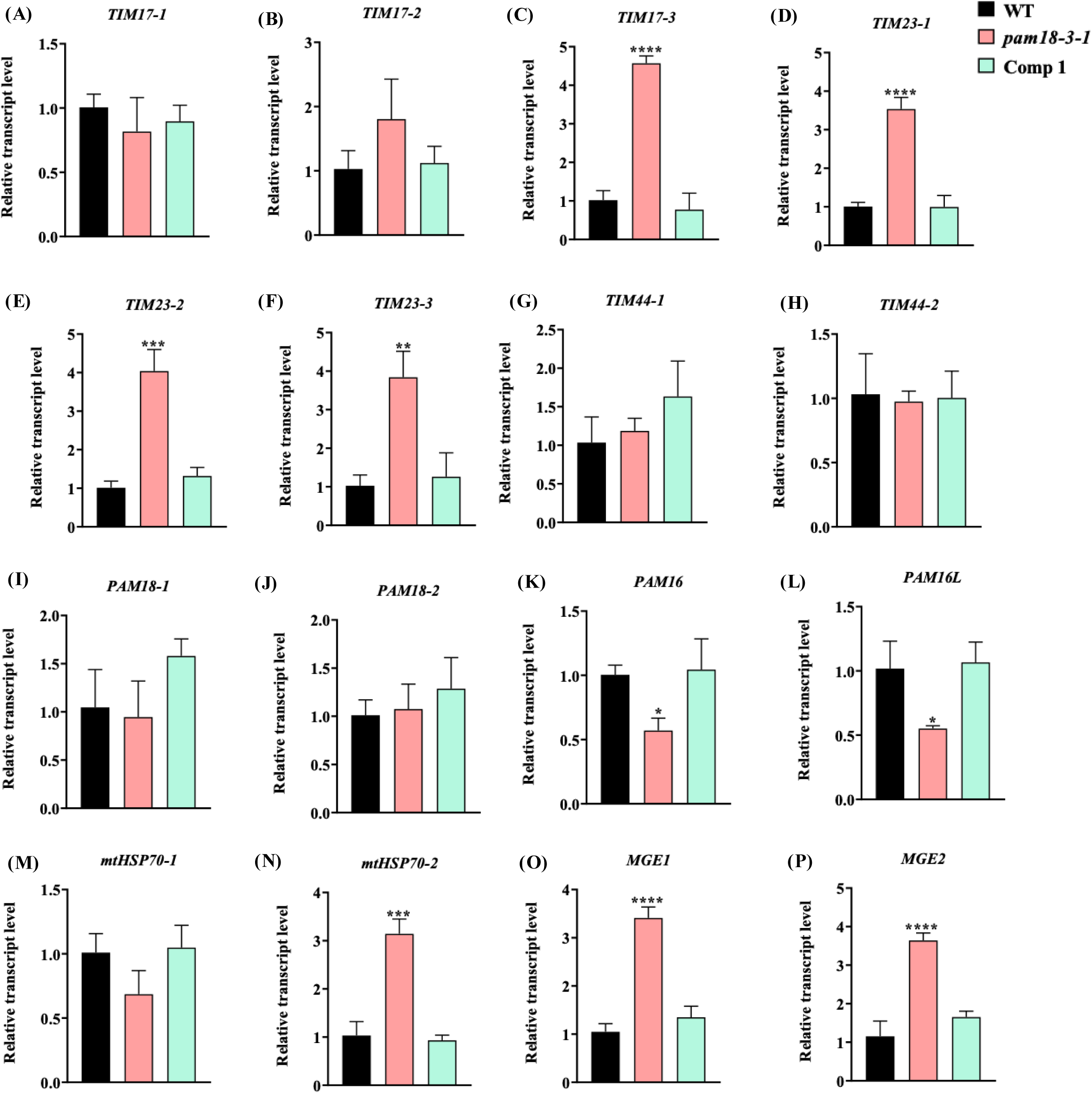
Transcript profiling of TIM23 complex components in 10-day-old Arabidopsis seedlings. qRT–PCR analysis of TIM23 complex component transcripts in WT, *pam18-3-1*, and complemented (Comp 1) seedlings. **(A–P)** Transcript levels of **(A)** *TIM17-1,* **(B)** *TIM17-2,* **(C)** *TIM17-3,* **(D)** *TIM23-1,* **(E)** *TIM23-2,* **(F)** *TIM23-3,* **(G)** *TIM44-1,* **(H)** *TIM44-2,* **(I)** *PAM18-1,* **(J)** *PAM18-2,* **(K)** *PAM16,* **(L)** *PAM16L,* **(M)** *mtHSP70-1,* **(N)** *mtHSP70-2,* **(O)** *MGE1,* and **(P)** *MGE2*. Expression levels were normalized to *ACT2* and *GAPDH*. Data represent the mean ± SD from three biological replicates, each with technical triplicates. Statistical analysis was conducted using one-way ANOVA with Tukey’s HSD test. Asterisks indicate significant differences compared to WT (* *p* < 0.05, ** *p* < 0.01, *** *p* < 0.001, **** *p* < 0.0001).

### Protein import via the TIM23 pathway is compromised in the *pam18-3* mutant

The TIM23 complex mediates the import of nearly 60% of the mitochondrial proteome, including all matrix-targeted proteins and a subset of inner mitochondrial membrane (IMM) proteins (Crameri et al., 2024; Ford et al., 2022). Considering the evolutionary conservation of PAM18 across eukaryotes, and the observed alterations in transcript levels of various TIM23 components in *pam18-3* mutants provide compelling evidence for a compromised mitochondrial protein import machinery. To further explore this, we quantified the steady-state abundance of three canonical TIM23 substrates: Isocitrate dehydrogenase (IDH), ATP synthase β-subunit (ATPβ), and Serine hydroxymethyltransferase 1 (SHMT1), in total cell lysates and purified mitochondrial fractions from wild-type, *pam18-3-1* mutants, and complemented (Comp 1) lines using quantitative immunoblotting. Coomassie Brilliant Blue (CBB) staining served as a loading control, and GAPDH was used as a cytosolic marker (Figure 7). In total cell lysates, the abundance of IDH, ATPβ, and SHMT1 was comparable in the three genotypes analyzed, indicating that overall expression of these proteins was not affected by the loss of *PAM18-3* (Figure 7). However, in the mitochondrial fractions, a marked reduction in the levels of all three substrates was observed in *pam18-3-1* relative to WT, suggesting a defect in mitochondrial import rather than protein synthesis. Quantification of the immunoblot bands revealed that the mitochondrial abundance of IDH was reduced to approximately 52%, ATPβ to 68%, and SHMT1 to 40% of WT levels (Figure 7B–D). Importantly, these defects were rescued in Comp 1, implying that the import deficiency was directly attributable to the loss of *PAM18-3* function (Figure 7B-D). These results demonstrate that *PAM18-3* is required for efficient protein translocation via the TIM23 complex, and its absence compromises the import of presequence containing nuclear-encoded mitochondrial proteins.

**Figure 7.**
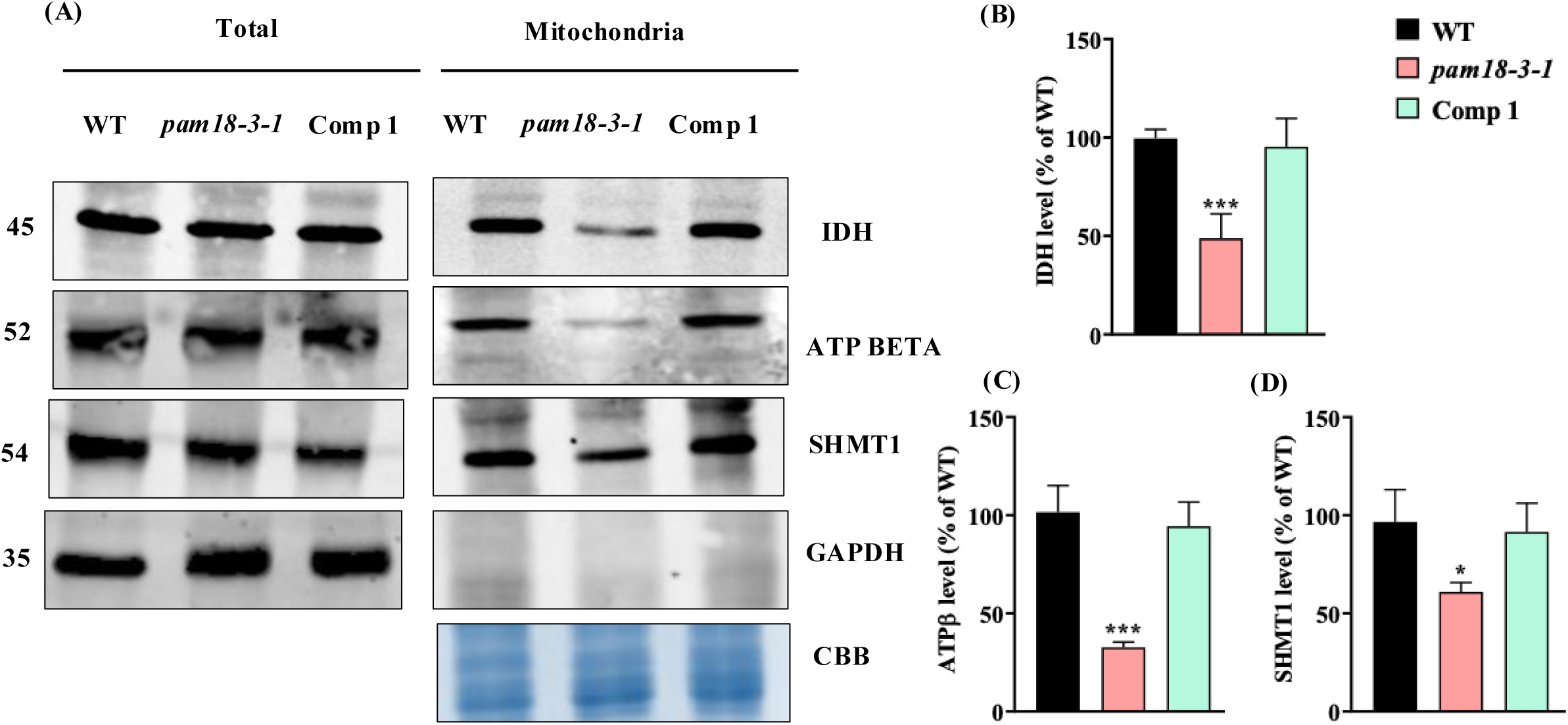
Immunoblot analysis of mitochondrial TIM23 pathway substrates in WT, *pam18-3-1*, and complemented (Comp 1). **(A)** Immunoblot analysis of total cell lysates (30 µg) and crude mitochondrial fractions (20 µg) from 10-day-old seedlings of WT, *pam18-3-1*, and complemented line (Comp 1), probed with antibodies against three mitochondrial proteins imported via the TIM23 complex: Isocitrate dehydrogenase (IDH), ATP synthase β-subunit (ATPβ), and Serine hydroxymethyltransferase 1 (SHMT1). Coomassie Brilliant Blue (CBB) staining served as a loading control, and GAPDH was used as a cytosolic marker. **(B–D)** Quantification of mitochondrial levels of IDH **(B)**, ATPβ **(C)**, and SHMT1 **(D)**, normalized to total protein (CBB) and expressed relative to WT, which was set to 100%. Bar graphs represent mean ± SD from three biological replicates. Asterisks indicate significant differences compared to WT (one-way ANOVA with Tukey’s HSD test; * *p* < 0.05, and *** *p* < 0.001).

### *pam18-3* mutant exhibits elevated ROS levels and induction of mitochondrial stress response pathways

Mitochondrial dysfunction often triggers the accumulation of reactive oxygen species (ROS) (Huang et al., 2016; Murphy, 2009). To determine whether the mitochondrial defects observed above perturb cellular redox homeostasis, ROS levels were analyzed in WT, *pam18-3-1*, and Comp1. 10-day-old seedlings were subjected to histochemical staining targeting two key ROS species: superoxide and hydrogen peroxide. Superoxide levels were visualized using nitroblue tetrazolium (NBT), which reacts with superoxide anions to form a blue formazan precipitate (Kumar et al., 2014). *pam18-3-1* seedlings displayed markedly stronger NBT staining than WT, indicating elevated superoxide levels (Figure 8A, B). Additionally, Hydrogen peroxide was detected using 3,3′-diaminobenzidine (DAB), which produces a brown precipitate in the presence of H₂O₂ (Daudi and O’Brien, 2012). Enhanced DAB staining in *pam18-3-1* seedlings further confirmed increased peroxide accumulation (Figure 8C, D). Quantitative analysis of staining intensities confirmed significantly enhanced NBT and DAB signals in *pam18-3-1* relative to WT, whereas the complemented line Comp1 restored ROS levels comparable to WT (Figure 8B, D). These results demonstrate that elevated ROS levels in *pam18-3-1* are directly associated with loss of *PAM18-3* function and its impact on mitochondrial integrity.

**Figure 8.**
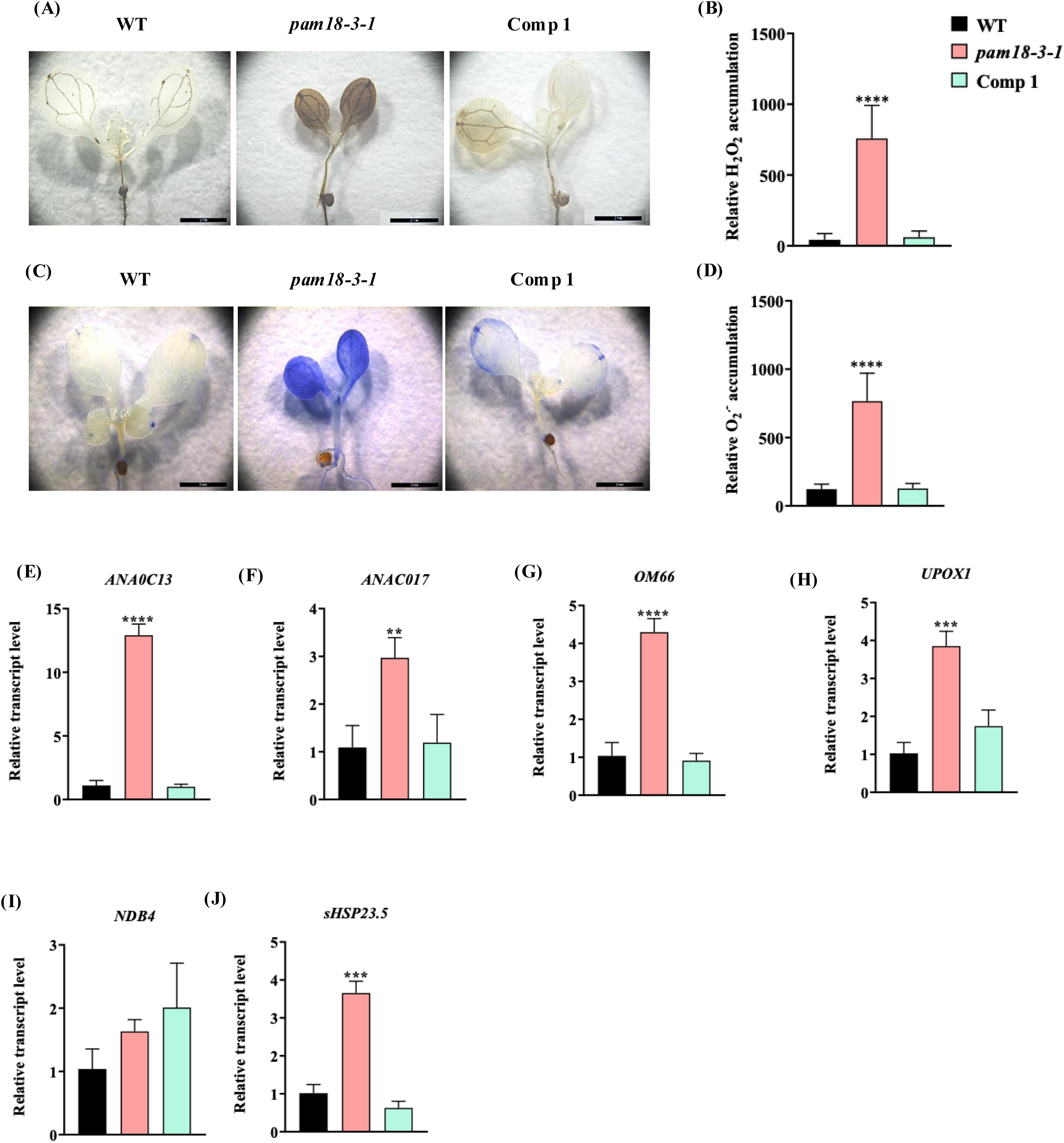
Accumulation of reactive oxygen species (ROS) and expression of mitochondrial dysfunction stimulon (MDS) genes in WT, *pam18-3-1*, and complemented (Comp 1) seedlings. **(A)** Detection of hydrogen peroxide (H₂O₂) accumulation by DAB staining in 10-day-old seedlings of WT, *pam18-3-1*, and Comp 1. **(B)** Quantification of H₂O₂ staining intensity using ImageJ. **(C)** Detection of superoxide (O₂•⁻) accumulation by NBT staining in WT, *pam18-3-1*, and Comp 1 seedlings. **(D)** Quantification of O₂•⁻ staining intensity using ImageJ. Scale bars in (A, C) = 2 mm. **(E–J)** Transcript levels of MDS marker genes assessed by qRT–PCR in 10-day-old WT, *pam18-3-1*, and Comp 1 seedlings: (E) *ANAC013*, (F) *ANAC017*, (G) *OM66*, (H) *UPOX1*, (I) *NDB4*, and (J) *sHSP23.5*. Expression levels were normalized to *ACT2* and *GAPDH*. Data represent the mean ± SD from three biological replicates, each with technical triplicates. Statistical analysis was conducted using one-way ANOVA followed by Tukey’s HSD test. Asterisks indicate significant differences compared to WT (\**p* < 0.05, \*\**p* < 0.01, \*\*\**p* < 0.001, **** *p* < 0.0001).

Accumulation of ROS in plant cells acts as a critical signal of mitochondrial dysfunction, triggering retrograde signaling pathways that alter nuclear gene expression to mitigate stress and restore cellular homeostasis (Huang et al., 2016; Khan et al., 2024). The genes collectively known as the Mitochondrial Dysfunction Stimulon (MDS) are a central part of this response, and their transcriptional activation is recognized as a hallmark of mitochondrial stress induced by elevated ROS levels (Meng et al., 2019; Ng et al., 2013). To determine whether this mitochondrial stress response is up-regulated in *pam18-3-1* mutants, transcript levels of several well-characterized MDS genes, including *AOX1A*, *MGE1*, *UPOX1*, *OM66*, *HSP23.5*, and the NAC transcription factors *ANAC013* and *ANAC017*, were quantified by qRT-PCR. All tested genes, with the exception of *NDB4*, exhibited significant upregulation in *pam18-3-1* relative to the WT, consistent with activation of ROS-mediated retrograde signalling (Figure 8E-J). Notably, transcript levels of these MDS genes in the Comp 1 were comparable to WT (Figure 8E-J), supporting the conclusion that *PAM18-3* loss is responsible for triggering mitochondrial stress responses through elevated ROS signaling.

## Discussion

Mitochondrial biogenesis and function rely extensively on the import of nuclear-encoded proteins through a dedicated translocase machinery. In plants, the TIM23 complex serves as a major conduit for preproteins targeted to the mitochondrial matrix or inner membrane. Our findings identify *PAM18-3*, a J-domain protein as a key component of TIM23-mediated protein import, and demonstrate its critical role in maintaining mitochondrial homeostasis thereby affecting plant growth and development in *Arabidopsis thaliana*.

Although Arabidopsis encodes three PAM18 paralogs with high sequence similarity and conserved domain architecture, our expression analysis revealed *PAM18-3* to be the most abundantly and ubiquitously expressed. Importantly, all three paralogs were capable of stimulating the ATPase activity of mtHSP70-1 to comparable levels, approximately 2.5 fold (Supplementary Figure S5; Supplementary Method), indicating that each retains the core biochemical function of J-domain proteins. Nevertheless, the pronounced developmental and mitochondrial defects observed exclusively in *pam18-3* mutants, despite the presence of *PAM18-1* and *PAM18-2*, suggest that its elevated and widespread expression is crucial for normal plant growth and mitochondrial function, pointing to functional specialization within this gene family. This pattern of specialization is consistent with observations in other eukaryotes. In *Saccharomyces cerevisiae*, the single PAM18 ortholog is indispensable for mitochondrial protein import and cell viability (D’Silva et al., 2003; Truscott et al., 2003). Similarly, in mammals, the PAM18 ortholog DNAJC19 plays a critical role, and its deficiency leads to DCMA (dilated cardiomyopathy with ataxia) syndrome, underscoring the essential and non-redundant function of this J-domain protein in mitochondrial homeostasis (Davey et al., 2006; Calvo et al., 2012). Notably, mammals also possess a related paralog, DNAJC15, which, although structurally similar, has distinct regulatory roles and tissue-specific functions (Radkevich et al., 2021; Cikánová et al., 2012), exemplifying how gene duplication can drive sub-functionalization or neo-functionalization within the PAM machinery.

In line with this functional specialization, loss-of-function mutants of *PAM18-3* exhibit pleiotropic developmental phenotypes, including severe growth retardation, altered rosette morphology, reduced seed set, and increased seed abortion. These defects phenocopy phenotypes observed in mutants of other TIM23-associated components (e.g., *mtHSP70-1*, *TIM17-1*, *PAM16*, *TIM23-1/3*), further highlighting the essentiality of the import apparatus for embryogenesis and post-embryonic growth (Huang et al., 2013; Li et al., 2021; Murcha et al., 2014; Wang et al., 2014; Wei et al., 2019). These pleiotropic defects reflect the broader importance of mitochondria in coordinating energy production, metabolic signaling, and stress responses throughout plant development.

Given the established role of PAM18 as an integral component of the presequence translocase-associated motor (PAM) complex in yeast, and its functional conservation in more complex eukaryotes, it is likely that *PAM18-3* is similarly essential for the efficient import of presequence-bearing proteins into the mitochondrial matrix in Arabidopsis. Consistent with this role, a reduction in the mitochondrial accumulation of several canonical TIM23-dependent substrates was detected in *pam18-3* mutants, including isocitrate dehydrogenase (IDH), ATP synthase β-subunit (ATPβ) and serine hydroxymethyltransferase 1 (SHMT1). This defect in protein import was accompanied by a pronounced decrease in mitochondrial membrane potential (Δψm) and compromised respiratory activity, both of which are key drivers of TIM23-mediated translocation. Given the reliance of preprotein import on a sustained proton motive force (Martin et al., 1991; Voos et al., 1999), the loss of Δψm in *pam18-3* likely aggravates the import deficiency by limiting the electrochemical gradient necessary for translocation thereby affecting respiratory activity. Together, these observations suggest a bidirectional relationship between mitochondrial protein import and bioenergetics, in which impaired import compromises respiratory efficiency, and reduced respiration further weakens import capacity, creating a potentially deleterious feedback loop that exacerbates mitochondrial dysfunction (Ferramosca, 2020). Consistent with this, transmission electron microscopy revealed swollen and morphologically irregular mitochondria, structural features commonly associated with mitochondrial import stress and loss of organellar integrity (Galloway and Yoon, 2013; Javadov et al., 2018). Together, these findings suggest that defects in protein import may act as the primary insult in *pam18-3* mutants, triggering a cascade of mitochondrial dysfunction including loss of ΔΨm and compromised ultrastructure.

On one hand, while the mitochondrial dysfunction in *pam18-3* mutants led to altered mitochondrial morphology and decreased ΔΨm, an unexpected increase in total oxygen consumption was noteworthy. This paradox is explained by a compensatory induction of the alternative oxidase (AOX) pathway, which accepts electrons from the ubiquinone pool and transfers them directly to oxygen, bypassing complexes III and IV (Saha et al., 2016; Vishwakarma et al., 2015). Activation of AOX is a conserved response to mitochondrial stress, often coordinated through ANAC transcription factors (Ng et al., 2013). Indeed, in *pam18-3* mutants, significant upregulation of *ANAC013*, *ANAC017*, *UPOX1*, *OM66*, *sHSP23.5*, and *NDB4* was observed, consistent with a robust engagement of the mitochondrial dysfunction stimulon (MDS). These data indicate that loss of *PAM18-3* triggers a multifaceted stress adaptation program involving both alternative respiration and stress-responsive gene expression. This included significant transcriptional upregulation of several TIM23 complex components, including *TIM17-3*, *TIM23-1*, *TIM23-2*, *TIM23-3* and *mtHSP70-2*, as well as chaperones and nucleotide exchange factors such as *MGE1* and *MGE2*. These transcriptional changes are characteristic of mitochondrial retrograde signaling, a process whereby functional deficits in mitochondria elicit nuclear gene expression adjustments to restore homeostasis. Interestingly, *PAM16* expression was suppressed, potentially reflecting a compensatory mechanism to alter co-chaperone dynamics in favor of enhanced mtHSP70 activity.

Despite AOX pathway induction, *pam18-3* mutants exhibited substantial accumulation of hydrogen peroxide (H₂O₂) and superoxide (O₂⁻·), as visualized by DAB and NBT staining, respectively. While AOX and MDS responses function to mitigate oxidative stress, persistent ROS accumulation suggests that the mitochondrial dysfunction exceeds the capacity of these protective systems. These results highlight a direct link between compromised mitochondrial protein import machinery and oxidative stress, in line with current models connecting mitochondrial dysfunction to redox imbalance and stress signaling in plants (Huang et al., 2016; Pan and Shadel, 2009). Elevated ROS in *pam18-3* may reinforce retrograde signaling via ROS-sensitive transcription factors, thereby contributing to the developmental and physiological defects (Huang et al., 2016).

Together, our findings underscore the intimate link between mitochondrial protein import, bioenergetics, redox balance, and plant development, positioning *PAM18-3* as a central regulator of mitochondrial function in *Arabidopsis thaliana* (Figure 9). While *PAM18-3* clearly fulfills a conserved role akin to yeast PAM18 and mammalian DNAJC19, as a core cochaperone of the TIM23/PAM system, potential additional roles beyond canonical protein translocation cannot be ruled out. Emerging evidence from yeast and mammalian systems implicates PAM18 homologs in diverse processes such as mitophagy, cardiolipin remodeling, and apoptotic regulation, suggesting broader involvement in mitochondrial quality control and signaling. Whether *PAM18-3* contributes to similar pathways in plants remains an open and intriguing question. Future investigations into its interaction partners, regulatory dynamics within the TIM23 complex, and possible roles in stress signaling and proteostasis will be crucial to fully elucidate its functions in plants.

**Figure 9.**
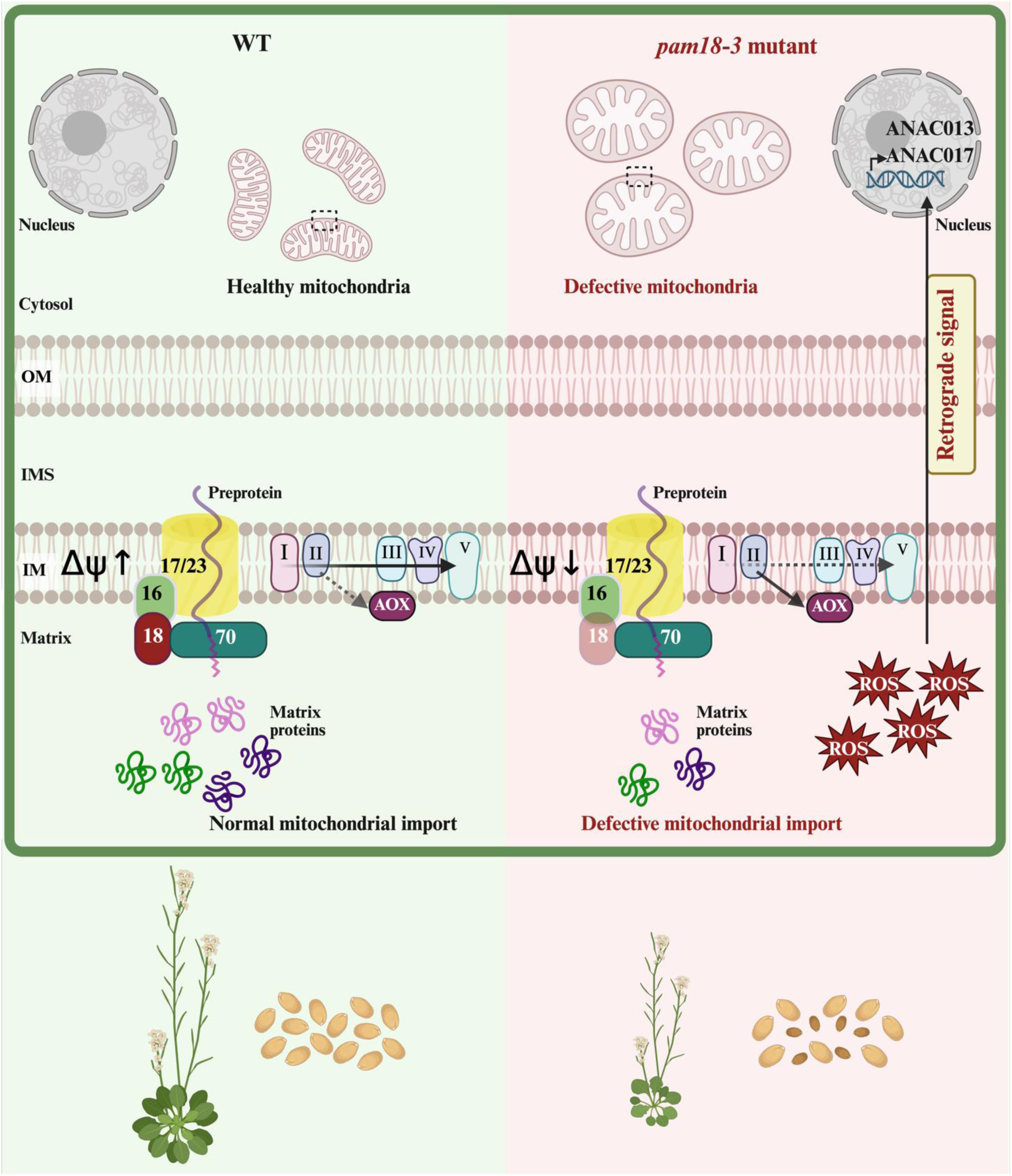
Proposed model for the role of *PAM18-3* in mitochondrial function and plant development. WT mitochondria maintain ΔΨ, support efficient protein import, and show major electron flow via the COX pathway (solid lines) with minor flow through AOX (dotted lines), sustaining normal growth and seed development. In the *pam18-3* mutant, mitochondria are swollen with reduced ΔΨ, impaired protein import, and altered respiration, where major electron flow shifts to AOX (solid lines) and minor flow persists through COX (dotted lines), leading to ROS accumulation, retrograde signalling, growth defects, and seed abortion. Key components of the import machinery are indicated: PAM18 (18), PAM16 (16), mtHSP70 (70), and the TIM17/23 translocase (17/23).

## Acknowledgements

The authors thank Dr. Elizabeth Vierling (Professor Emerita, Department of Biochemistry & Molecular Biology, University of Massachusetts Amherst) and Dr. Chetana Tamadaddi (Brandizzi Lab, Michigan State University, USA) for insightful scientific discussions. The authors also thank the Electron Microscopy Facility at Vienna BioCenter Core Facilities (VBCF), part of the Vienna BioCenter (VBC), Austria, for performing the TEM experiment. N.S. gratefully acknowledges the Council of Scientific and Industrial Research (CSIR) for fellowship support (09/1020(0197)/2020-EMR-I). Figures were created with BioRender.com (https://biorender.com).

## Author contributions

C.S. and Neha conceptualized the study, designed the experiments, and wrote the manuscript. C.S. supervised the research and provided laboratory facilities and funding. Neha performed the majority of the experiments and analyzed the data. S.S. contributed to phenotyping and mitochondrial protein assays.

---

**Figure S1.**
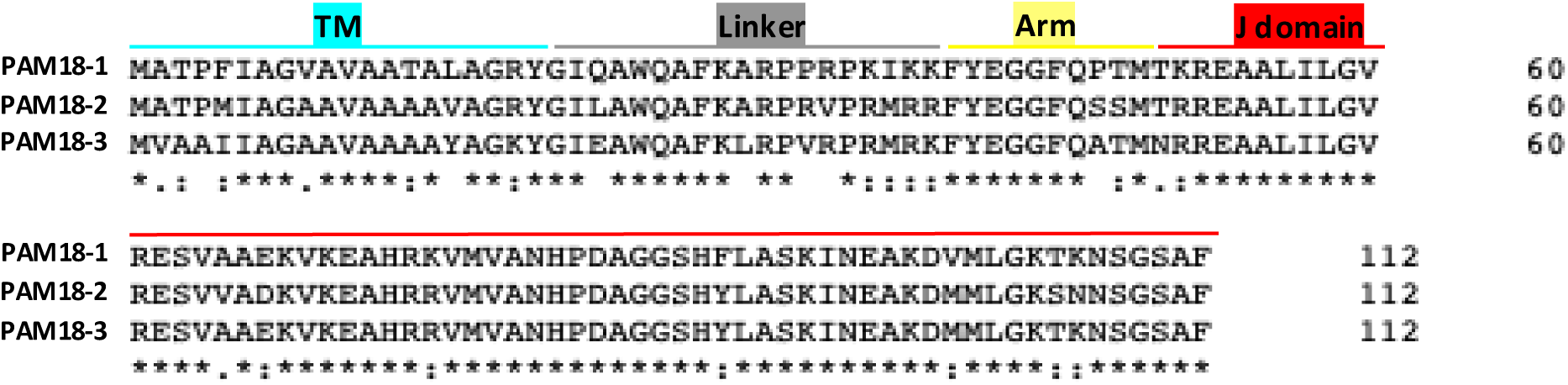
Multiple sequence alignment of Arabidopsis PAM18 proteins. Amino acid sequence alignment of Arabidopsis PAM18- 1, PAM18-2, and PAM18-3 proteins was performed using ClustalW. Identical residues are indicated by asterisks (*), conserved substitutions by colons (:), and semi-conserved substitutions by periods (.). Functional domains are highlighted: transmembrane helix (TM, cyan), linker region (grey), arm domain (yellow), and J-domain (red).

**Figure S2.**
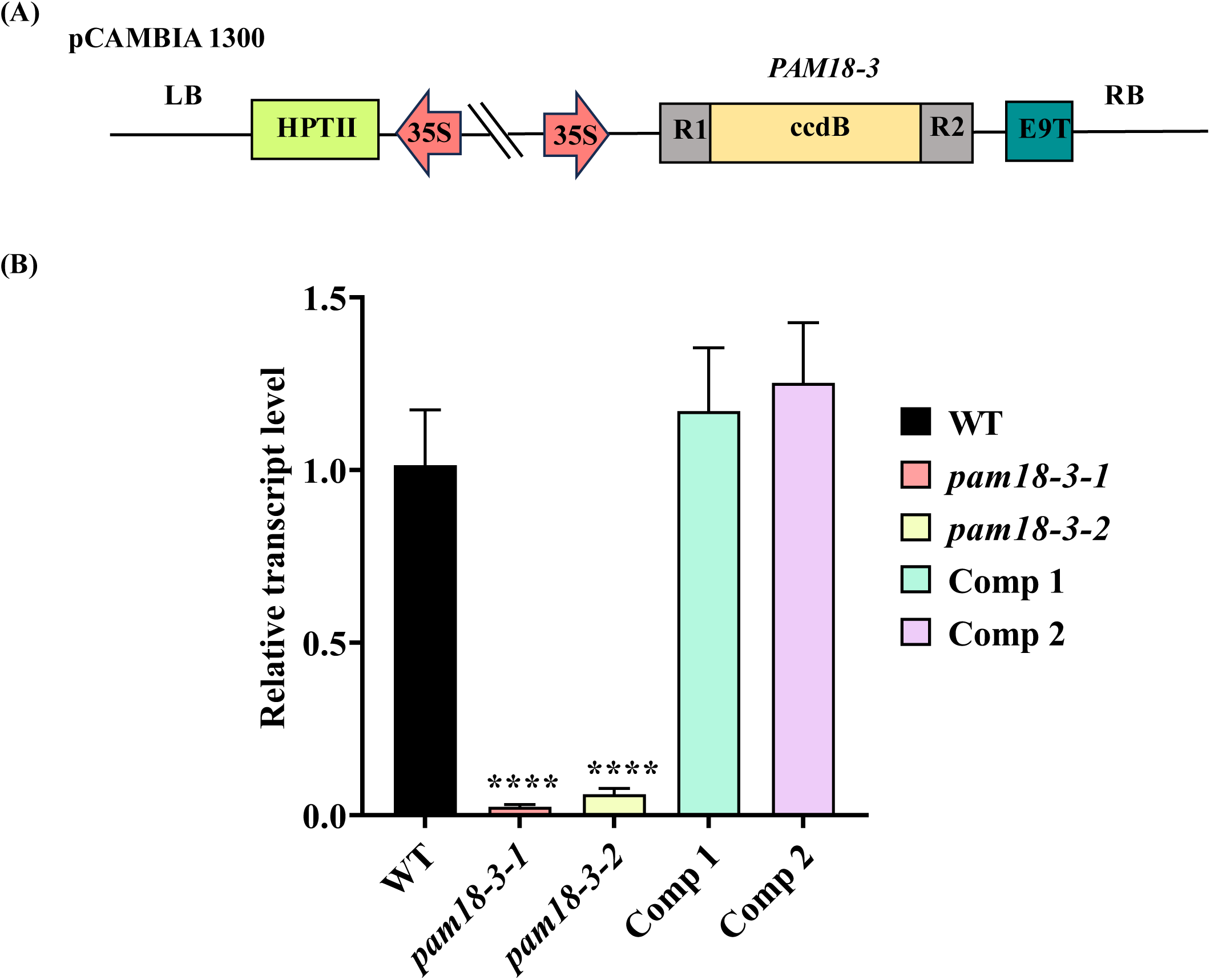
Characterization of *pam18-3* mutant and complemented lines for phenotypic analysis. **(A)** Schematic representation of the pCAMBIA1300 binary vector used for generating complemented lines of *AtPAM18-3*. **(B)** Relative expression levels of *AtPAM18-3* in 10-day-old seedlings of WT, two independent mutant alleles (*pam18-3-1* and *pam18-3-2*), and two independent complemented lines (Comp 1 and Comp 2). Transcript levels were normalized to *ACT2* and *GAPDH*. Data represent the mean ± SE of three biological replicates. Asterisks indicate significant differences compared to WT (one-way ANOVA with Tukey’s HSD test; (\*\*\*\**p* < 0.0001).

**Figure S3.**
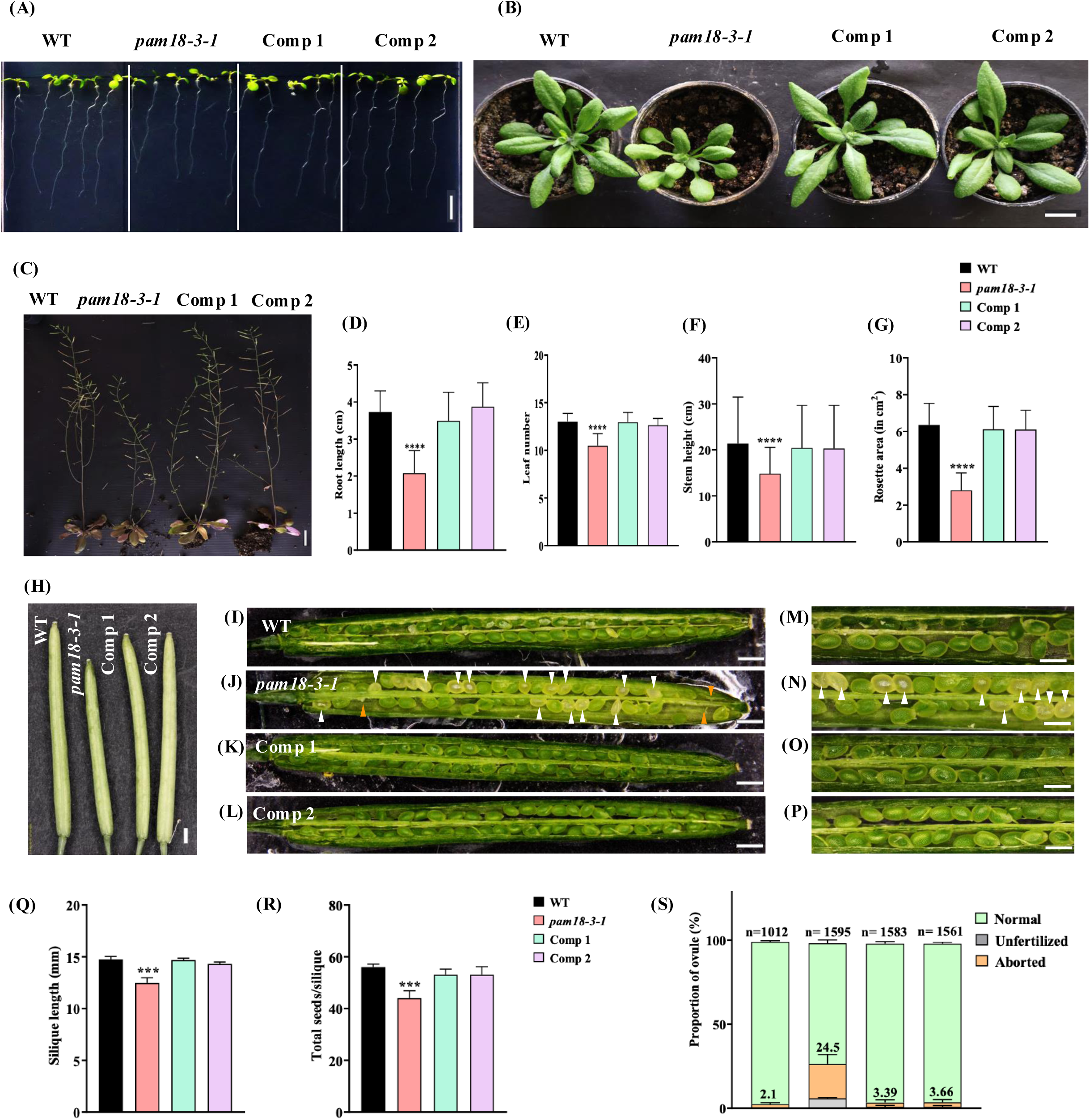
Phenotypic characterization of *pam18-3-1* mutant and complemented lines (Comp1 and Comp 2). **(A–C)** Representative growth phenotypes of WT, *pam18-3-1*, and two complemented lines (Comp 1, Comp 2): 10-day-old seedlings **(A)**, 21- day-old rosettes **(B)**, and 45-day-old flowering plants **(C)**. **(D–G)** Quantitative analysis of root length (D), total leaf number **(E)**, stem height **(F)**, and rosette area **(G)**. Data represent mean ± SD from three independent experiments (*n* = 30 plants per genotype). Statistical significance was determined using one-way ANOVA followed by Tukey’s multiple comparison test (****p < 0.0001). Scale bars: A, B = 1 cm; C = 2 cm. **(H)** Representative siliques at 8 days post-anthesis (DPA). **(I–P)** Seed development in 8-DPA siliques: WT **(I, M)**, *pam18-3-1* **(J, N)**, Comp 1 **(K, O)**, and Comp 2 **(L, P)**. Aborted ovules are indicated by white arrowheads, unfertilized ovules by orange arrowheads. Scale bars: I, J, L, N = 1 mm; K, M, O, P = 0.5 mm. **(Q–S)** Quantification of silique length **(Q)**, total seeds per silique **(R)**, and proportion of normal, unfertilized, and aborted ovules per silique **(S)**. Data represent mean ± SD from three biological replicates (*n* = 10 siliques from 10 individual plants per replicate). Statistical analysis was performed using one-way ANOVA with Tukey’s HSD test (\*\*\**p* < 0.001).

**Figure S4.**
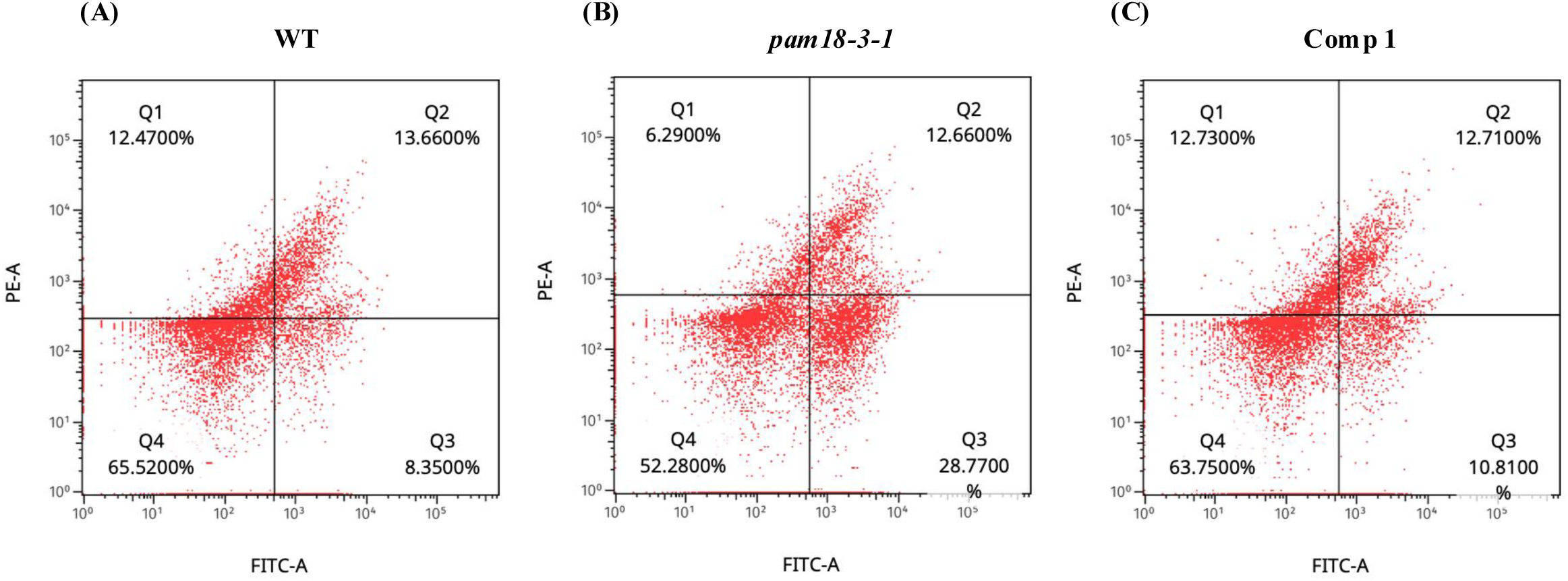
Flow cytometric analysis of WT, *pam18-3-1* mutant, and complemented (Comp 1) lines. Representative dot plots showing FITC-A fluorescence (x-axis) versus PE-A fluorescence (y-axis) for WT, *pam18-3-1* mutant, and Comp 1 lines. Each plot is divided into four quadrants: Q1 (PE-positive / FITC-negative), Q2 (PE-positive / FITC-positive; double-positive), Q3 (FITC-positive/ PE-negative), and Q4 (double-negative). Percentages within each quadrant represent the proportion of total events for that population. Data shown are representative of three independent biological replicates.

**Figure S5.**
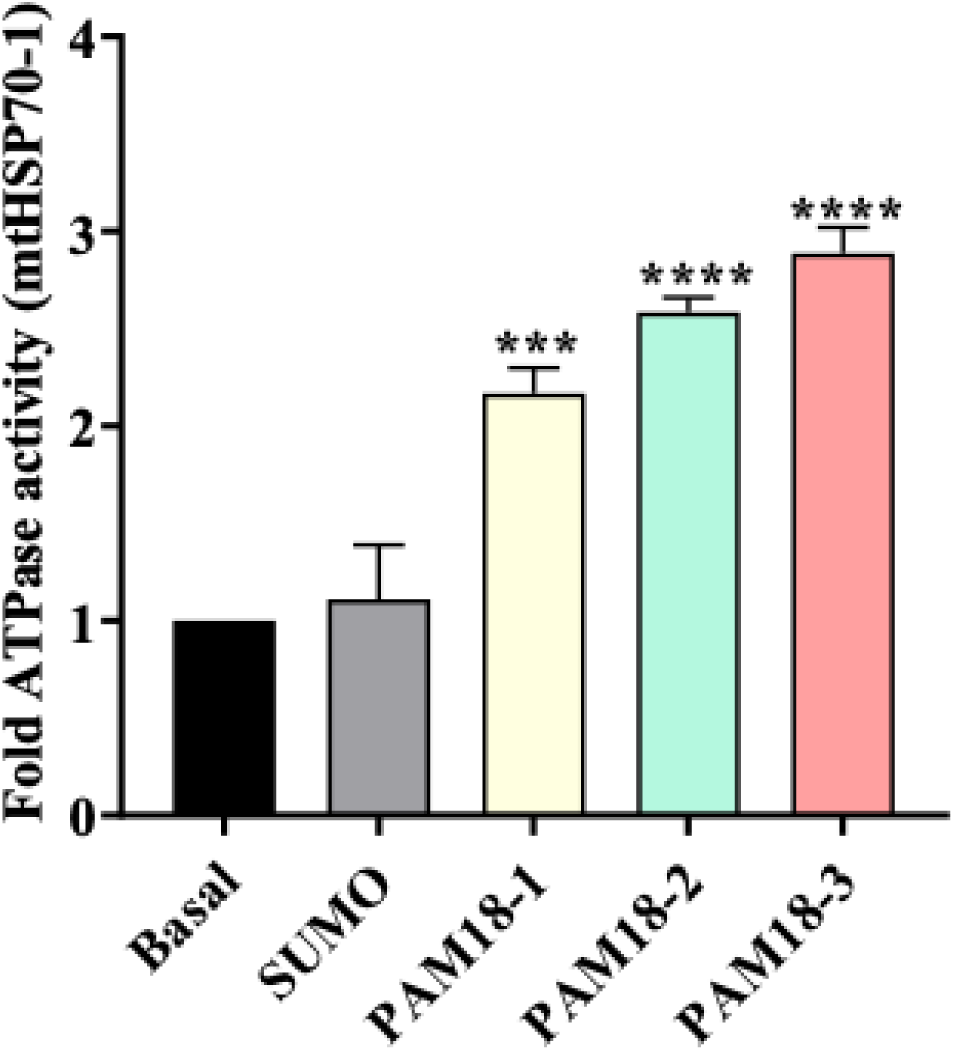
Relative stimulation of mtHSP70-1 ATPase activity by Arabidopsis PAM18 paralogs. ATPase activity was measured using a malachite green–based assay. mtHSP70-1 was incubated with a four-fold molar excess of SUMO-tagged (C-terminus) PAM18-1, PAM18-2, or PAM18-3, and ATP hydrolysis was quantified after 1 h at 23 °C. The basal ATPase activity of mtHSP70-1 alone was normalized to 1, and SUMO tag alone was included as a control. Data are shown as fold stimulation relative to mtHSP70-1. Error bars represent mean ± SEM (*n* = 3 independent assays). Statistical analysis was performed using one-way ANOVA with Tukey’s HSD test (\*\*\**p* < 0.001, and \*\*\*\**p* < 0.0001).

