## Supplementary TableS1 for "PAM18-3, a J-domain protein, maintains mitochondrial integrity and plant growth and development in *Arabidopsis thaliana*"

**Table S1.** List of primers used in this research

| **Primer Name** **Sequence (from 5´ to 3´)**  **Forward Reverse** | | |
| --- | --- | --- |
| **Promoter:PAM18-1/2/3:GUS primers** | | |
| Promoter PAM18-1 | GGGGACAAGTTTGTACAAAAAAGCAGGCTGGTCTCTCTTCCTCCACCTTCAC | GGGGACCACTTTGTACAAGAAAGCTGGGTGTCAGAAATCTAGGATTCTTCG |
| Promoter PAM18-2 | GGGGACAAGTTTGTACAAAAAAGCAGGCTGGCAAAACTTGAAGAGCTTCTCG | GGGGACCACTTTGTACAAGAAAGCTGGGTGTGTTGTAGAGGATTTGAGCTA |
| Promoter PAM18-3 | GGGGACAAGTTTGTACAAAAAAGCAGGCTGGATGTTCATGGATCCAAGCAAC | GGGGACCACTTTGTACAAGAAAGCTGGGTGTATTCAGCTAAGTAGTTGTTC |
| **Genotyping primers** | | |
| SALK_094163 | LP: CAACTCCCTCAAGGCTAAACC | RP: CTTCGATTGTCGAGTTTCTCG |
| SALK_091892C | LP: TGAGTGCCGTAAGGTTTATGG | RP: CTGGCTATACACAGGCTACGC |
| **Complementation primers (Gateway cloning)** | | |
| *PAM18-3* | GGGGACAAGTTTGTACAAAAAAGCAGGCTGGATGGCTACGCCAATGATTGCAGG | GGGGACCACTTTGTACAAGAAAGCTGGGTGTCAAAAGGCAGAACCGCTGTTGTTGG |
| **RT primers** | | |
| *PAM18-1* | GCAAGCATTCAAGGCAAGGC | TGATCTTAGAGGCTAGGAAATGG |
| *PAM18-2* | GGAATAGAGGCATGGCAAGC | CGCAAGGTAATGGCTTCCTC |
| *PAM18-3* | GGAGAGTGTTGTGGCAGATAAGG | ACCGCTGTTGTTGGATTTTC |
| *ANAC017* | CATCGGCTTTGTGGGCATTA | CACAAAGCACCCACGATTGA |
| *ANAC013* | GTGGTGGTCGTTTCAGGTTT | CTCCCTGAACCTCCCATTGT |
| *OM66* | GCGTGGACCTCTGTTACTCT | GCGGTCAACTCTAGGTCGTA |
| *UPOX1* | CAAACCTCAAGGATCACATGGATGA | GCCTTGGAGAAGCTCCCGAATATCT |
| *NDB4* | TCAAGTCTTAAGGGCACAACA | CGGAAGAGAGAGGCTCTCTCG |
| *sHSP23.5* | TCAAACCGACATGTTTCTCG | AAGCTTCTCGTTGGAGTAAACG |
| *MGE1* | ACGCAGTGTTCCAAGTCCCA | TCTTTGCCGCCTTCTTGATT |
| *MGE2* | GTATTTAAGAAATTTGGTATGG | TTAAGCATCAGACTCTTTCTTTTC |
| *TIM44-1* | GAAGTCTCAGTTTCTGTGAC | TTAAATGAGAGCTTGAACACC |
| *TIM44-2* | GTTGATATTCAAGAGACAAAG | TTAAATGAGAGCTTGAACACC |
| *mtHSP70-1* | CAGCTGAGATTGCCTCTGAG | TCACTTCCTTGAACCACTGG |
| *mtHSP70-2* | CAGAGAGAAGATCCCAAGTG | TCACTTTTTCACTTCCTCGT |
| *TIM17-1* | CATGGAGGATCATGCAGCTA | GTAGGCACAGGAGGAGC |
| *TIM17-2* | CATGGAGGATCATGCAGCTA | GCAGCACCAGCAATGATAGA |
| *TIM17-3* | TGAGAGGGGCATACAATTCC | CAACAACCCCTTTTCGGATA |
| *TIM23-1* | GTGCTTCAGCTGGGATCTTC | CAACACTGGTCCAAACATCG |
| *TIM23-2* | TTGGGGTAATCGGATTGGTA | CAACACTGGTCCAAACATCG |
| *TIM23-3* | CCGTAGGGTTGATGTTTGCT | ATTGGAACGAATCGCTTGAC |
| *AOX1A* | GTTTCGTCTCACGAGGCTTTAT | GGTGGATTCGTTCTCTGTTTTC |
| *AOX1B* | CAAGCTAATGGAAACTGCTGTG | CATCTTGCTGAAAACTCTCACG |
| *AOX1C* | ATTACTCCGTCGCTCTCTCCTT | CTTCACGCCCCAATAACTAACT |
| *AOX1D* | GGATTCAGGGGACATCTCATTA | CTGGCTGGTTATTCCCACTTAC |
| *AOX2* | CATGTTCGTGAGTTCTGTTTCC | ACCCATCCACCTCAAGTTAAAA |
| *ACTIN2* | TATCGCTGACCGTATGAGCAAAG | TGGACCTGCCTCATCATACTCG |
| *GAPDH* | TTGGTGACAACAGGTCAAGCA | AAACTTGTCGCTCAATGCAATC |
