## Supplementary method for "PAM18-3, a J-domain protein, maintains mitochondrial integrity and plant growth and development in *Arabidopsis thaliana*"

**ATPase assay**

Functional fragments of PAM18 (aa 24–112) and mtHSP70-1 (aa 45–683), both lacking their mitochondrial targeting sequences, were cloned into the pSMT3 vector to generate N-terminal SUMO–Hexa-Histidine fusions.  Recombinant proteins were expressed in *E. coli* Rosetta cells and purified to homogeneity using Ni-NTA agarose beads (Qiagen) according to the manufacturer’s protocol. ATPase activity of purified mtHSP70-1 was determined using a malachite green–based colorimetric assay, essentially as described by Chang et al. (2008) with minor modifications. Purified mtHSP70-1 (1 µM) was incubated either alone or with a four-fold molar excess (4 µM) of C-terminal SUMO-tagged PAM18-1, PAM18-2, or PAM18-3 in assay buffer containing 100 mM Tris-HCl (pH 7.5), 20 mM KCl, 6 mM MgCl₂, and 0.017% Triton X-100, in a final reaction volume of 200 µL. A SUMO-only control (4 µM) was included to exclude non-specific effects of the tag. Reactions were initiated by addition of 1 mM and incubated for 1 h at 23 °C. ATP hydrolysis was detected by addition of malachite green reagent, and absorbance was measured at 620 nm using a microplate reader. Inorganic phosphate concentration was calculated from a KH₂PO₄ standard curve, and background absorbance from buffer-only blanks was subtracted. ATPase activity was expressed as pmol ATP hydrolyzed per µg mtHSP70-1 per min, and values were normalized to the basal activity of mtHSP70-1 alone (set as 1-fold).
